# *SpatialBench*: Comparative cross-platform benchmarking of high-resolution spatial transcriptomics using matched mouse lymphoid tissue

**DOI:** 10.64898/2026.04.29.721531

**Authors:** Ashleigh Solano, Raymond K. H. Yip, Changqing Wang, Daniela Amann-Zalcenstein, Pradeep Rajasekhar, Ishrat Zaman, Allan Motyer, Marek Cmero, Yang Xu, Yining Pan, Casey J. A. Anttila, Stephanie I. Studniberg, Peter F. Hickey, Layla Wang, Callum J. Sargeant, Ling Ling, Yunshun Chen, Ruvimbo D. Mishi, Lisa J. Ioannidis, Kim L. Good-Jacobson, Hamish W. King, Kelly L. Rogers, Diana S. Hansen, Rory Bowden, Matthew E. Ritchie

**Author notes:** These authors contributed equally as co-first (Ashleigh Solano, Raymond K. H. Yip, Changqing Wang, Daniela Amann-Zalcenstein). These authors contributed equally as co-last (Kelly L. Rogers, Diana S. Hansen, Rory Bowden, Matthew E. Ritchie).

## Abstract

Spatial transcriptomics (ST) has rapidly expanded with the introduction of multiple high-resolution platforms, yet cross-platform benchmarking remains limited and largely focused on technical performance. Here we present *SpatialBench*, a matched multi-platform resource comprising Visium HD, Xenium and MERSCOPE data together with single-cell and single-nucleus references from a malaria-challenged wild-type and B cell-specific *Tbx21* knockout mouse spleen model. In this system, loss of T-bet in B cells disrupts germinal center (GC) polarization and antibody maturation, providing a biologically grounded benchmark for technology comparison. We leveraged this system to systematically evaluate ST platform performance using technical and biological readouts. Across platforms, immune organization and *Tbx21*-associated programs were consistently recovered, indicating robustness of major biological signals. Platforms instead differed in the level of biological resolution accessible. Visium HD enabled transcriptome-scale GC characterization and, together with Xenium, resolved dark and light zone organization, whereas GC zonation was not resolved in MERSCOPE, consistent with differences in transcript detection sensitivity. *SpatialBench* provides a biologically defined reference dataset for evaluation of ST technologies, method development, computational benchmarking, and studies of GC spatial organization in lymphoid tissue.

## Introduction

Spatial transcriptomics (ST) links gene expression to tissue architecture, offering mechanistic insight into processes where spatial organization underpins function^(1–3)^. This capability has transformed studies in development^(4,5)^, cancer^(6–9)^, and immunity^(10,11)^, yet rigorous cross-platform evaluation remains limited. With multiple commercial platforms now widely available, there is a pressing need for independent benchmarking resources that support fair technology comparison, reproducible analysis, and guide future method development.

To meet this need, it is essential to consider the current landscape of ST platforms, which divide broadly into imaging-based (iST) and sequencing-based (sST) approaches^(12,13)^. Leading iST platforms include Xenium (10x Genomics)^(14)^, MERSCOPE (Vizgen, based on Multiplexed Error-Robust Fluorescence In Situ Hybridization)^(15,16)^, and CosMx (NanoString, now Bruker Spatial Biology)^(17,18)^, which use iterative *in situ* hybridization and imaging to localize hundreds to thousands of transcripts with high spatial precision and single-molecule resolution. Panels are customizable but remain limited in breadth compared with sST assays. On the other hand, iST enables simultaneous RNA and protein measurements with tissue morphology, a capability not yet established for sST platforms. By contrast, sST methods capture transcripts on spatially barcoded arrays to provide near transcriptome-wide coverage, though historically at lower spatial resolution than imaging-based approaches. Early sST assays including Slide-seq (∼ 10 µm beads)^(19)^, HDST (∼ 2 µm barcoded arrays)^(20)^, and Stereo-seq (sub-micron DNA nanoball arrays)^(21)^ demonstrated that cellular and even subcellular resolution was technically possible, although low capture efficiency, protocol complexity, and limited commercial accessibility restricted widespread use. The original Visium (10x Genomics) became the most widely adopted sST platform^(13)^, offering transcriptome-wide coverage while aggregating expression from multiple cells per 55 µm resolution that aggregates expression from multiple cells per spot^(22,23)^. The recent introduction of Visium HD, with a 2 µm pitch and an effective bin size of 8 µm, now delivers sequencing-based profiling at near-cellular resolution in a commercially accessible format^(24)^, narrowing the gap with iST. These developments further emphasize the need for systematic benchmarking to define trade-offs and establish where each technology is most informative.

Recent benchmarking studies have evaluated both sST and iST technologies, revealing consistent platform trade-offs but also highlighting gaps in tissue diversity, experimental design, and cross-platform comparability. Sequencing-focused work by You et al.^(25)^ compared 11 spatial methods, including Visium, Slide-seq, HDST, and Stereo-seq, across multiple mouse tissues, demonstrating substantial variation in capture efficiency, transcript diffusion, and effective spatial resolution, and establishing the *cadasSTre* reference dataset for reproducible cross-platform analysis. Imaging-based bench-marking studies have expanded in parallel. Analyses in brain and tumor models showed that iST platforms can resolve anatomical features with higher spatial precision than sST approaches, but exhibit variability in detection efficiency, cell type annotation, and susceptibility to transcript misassignment^(26,27)^. However, these comparisons largely relied on independent tissue samples, limiting controlled platform comparisons. Matched-tissue comparisons have begun to address this limitation. For example, Cook et al. ^(28)^ reported higher sensitivity and dynamic range for Xenium compared to CosMx in prostate adenocarcinoma, but stronger default segmentation performance for CosMx. Extending this approach, Cervilla et al. ^(29)^ compared five platforms across six cancer types and reported improved data quality for Visium CytAssist relative to manual Visium workflows, higher specificity for Xenium compared to CosMx, and identified Visium HD as uniquely combining transcriptome-wide coverage with near-cellular resolution. Broader tissue surveys have further characterized platform-dependent performance differences. Wang et al. ^(30)^ analyzed 17 tumor and 16 normal tissue types, including lymphoid tissues such as spleen and tonsil, using Xenium, MERSCOPE, and CosMx, highlighting trade-offs in sensitivity, specificity, and biological resolution. Additional studies demonstrated platform differences in transcript assignment accuracy and cell boundary delineation^(31)^, consistent with best-practice analyses showing that segmentation performance and quality control requirements vary substantially across tissue contexts^(32)^. Recent multi-platform comparisons have emphasized the importance of matched multi-omic references. Ren et al. ^(33)^ bench-marked four subcellular-resolution platforms (Stereo-seq, Visium HD, CosMx and Xenium) and released the SPATCH resource integrating ST with CODEX protein imaging^(34)^ and scRNA-seq as a reference for computational benchmarking.

Together, these studies establish consistent platform trade-offs, with iST platforms excelling in spatial resolution and segmentation fidelity, while sST provides broader transcriptome coverage at the cost of lower detection efficiency and increased technical noise, increasing reliance on computational methods to resolve cell types and spatial domains. Nevertheless, three key limitations remain. First, most benchmarks focus on brain and solid tumor tissues^(26,28,29,33)^, whereas immune lymphoid organs remain underrepresented despite providing a particularly stringent benchmarking context. Lymphoid tissues combine well-defined spatial microenvironments, such as germinal center (GC) zonation, with highly dynamic immune cell states, requiring platforms to resolve both structured architecture and subtle spatial transitions. Where included, profiling has typically been limited to small cores within multi-tissue arrays using panels not designed for immune biology, precluding systematic evaluation of lymphoid architecture and immune spatial organization^(30)^. Second, biologically defined perturbations that provide ground truth are rarely incorporated. Third, even where multi-platform comparisons exist, studies often rely on FFPE tumor material or differing gene panels, limiting direct cross-platform comparability despite recent efforts to develop matched multi-omic resources^(33)^. Addressing these gaps requires benchmarking resources grounded in biologically validated systems using matched tissues and harmonized measurements across technologies.

Here we present *SpatialBench*, a multi-platform benchmark of high-resolution ST technologies in immune tissue. This work builds on our earlier *SpatialBenchVisium* resource^(35)^, which established a standardized analysis framework for malaria-challenged spleens using the original Visium assay. The expanded dataset comprises 27 matched fresh-frozen spleen sections from wild-type, control and B cell–specific T-bet (*Tbx21*^fl*/*fl^ *Cd23*-Cre) knockout mice infected with malaria and drug-cured prior to tissue collection, profiled using Visium HD, Xenium and MERSCOPE. This model provides a well-defined immunological reference system in which GCs, key sites of antibody maturation, are spatially organized into proliferative dark and selection-associated light zones. Conditional deletion of T-bet in mature B cells produces reproducible alterations in GC transcriptional programs and zonation, providing a biological phenotype against which platform performance can be evaluated. This design enables systematic comparison of transcriptome coverage, sensitivity, background signal and cell-segmentation performance across platforms. Parallel differential expression analyses compare the recovery of known T-bet-dependent programs, while spatial analyses evaluate each platform’s ability to resolve GC zonation. Anchored in a controlled biological system, *SpatialBench* provides a standardized reference dataset for cross-platform evaluation and a resource for studies of GC spatial organization.

## Results

### *SpatialBench* design and dataset generation

To enable systematic benchmarking of ST platforms, we designed *SpatialBench* around three high-resolution technologies, with Visium HD (10x Genomics) representing sST and MERSCOPE (Vizgen) and Xenium (10x Genomics) representing iST approaches (Fig. 1A). Fresh-frozen mouse spleens were selected because the spleen contains diverse immune cell subsets arranged in distinct microanatomical units, in which conditional deletion of T-bet in mature B cells disrupts dark and light zone organization, providing a ‘ground truth’ perturbation for technology comparison (Fig. 1B). Using this model, we generated 27 spatial datasets (4 Visium HD, 9 MER-SCOPE, 14 Xenium, including 9 unimodal and 5 multimodal staining) together with matched single-cell and single-nuclei references from 10x Genomics 3’ GEX and FLEX profiling of wild-type spleens (reverse-transcription (RT) and probe-based transcriptome wide assays, respectively) (Fig. 1B–D, Supplemental Table. 1). MERSCOPE and Xenium were profiled with 91- and 100-gene panels, respectively, while Visium HD captured 19,059 genes. A shared set of 90 genes provided a minimal, common basis for cross-platform comparison (Fig. 1D, Supplemental Table 2).

**Fig. 1.**
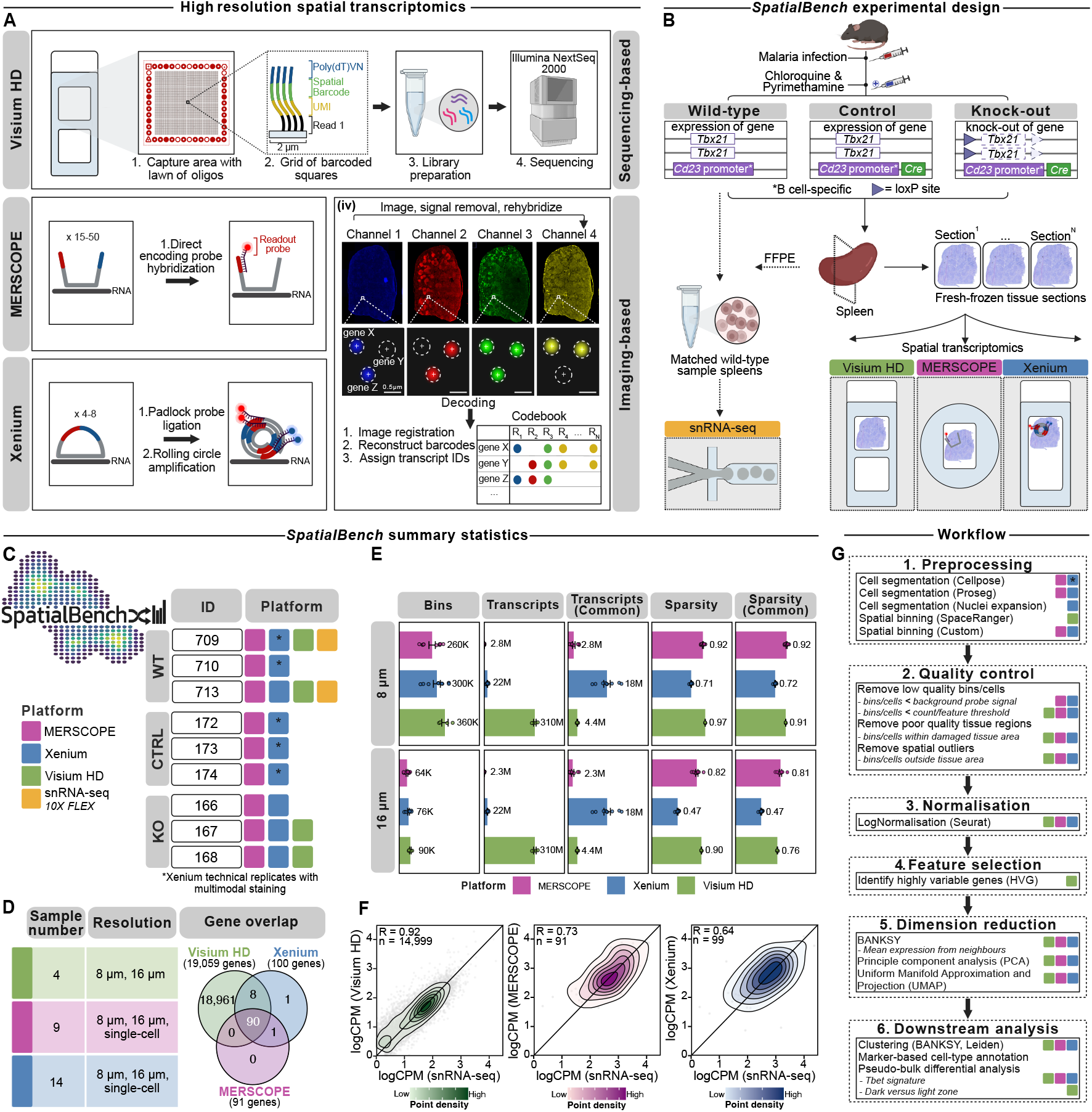
Overview of the *SpatialBench* study design for high-resolution spatial transcriptomics. **A**. Schematic overview of the three spatial transcriptomics (ST) platforms used in this study, comprising sequencing-based (sST) and imaging-based (iST) approaches. Visium HD (10x Genomics) is a sST platform that captures spatially barcoded polyadenylated transcripts, followed by cDNA library preparation and sequencing. MERSCOPE (Vizgen) and Xenium (10x Genomics) are iST platforms for *in situ* transcript detection using multiplexed fluorescence imaging, with MERSCOPE relying on direct fluorescent probe hybridization and Xenium on padlock probes with rolling circle amplification, followed by iterative imaging, signal removal, and rehybridization cycles for transcript decoding. **B**. Experimental design of the *SpatialBench* study using spleen sections from malaria infected and cured mice across wild-type (WT), control (CTRL) (*Tbx21*^+/+^ *Cd23*^Cre^), and B cell-specific conditional *Tbx21* knockout (KO) (*Tbx21*^fl/fl^ *Cd23*^Cre^) genotypes. Fresh-frozen spleen sections were profiled using Visium HD, MERSCOPE, and Xenium. Matched single-nuclei RNA seq (snRNA-seq) datasets were generated from dissociated formalin-fixed paraffin-embedded (FFPE) WT spleens. **C**. Sample composition of the *SpatialBench* dataset. Colors denote platform identity, with MERSCOPE (magenta), Xenium (blue), Visium HD (green), and snRNA-seq (yellow). Asterisk denote Xenium technical replicates with multimodal staining. **D**. Summary of sample number, spatial resolution, and Venn diagram displaying shared genes between panels of the ST platforms. **E**. Dataset-level statistics for binned ST data, with 8 µm and 16 µm bins shown in the top and bottom rows, respectively. Columns show total bins, total transcript counts, transcript counts for the subset of 90 shared genes, sparsity, and sparsity for the shared genes across platforms. Points indicate individual samples, with bars and error bars denoting the mean and standard error of the mean (SEM). **F**. Global Pearson correlation of gene expression between snRNA-seq and ST platforms. Density contour plots show gene-level correlations for Visium HD (green, left), MERSCOPE (magenta, middle), and Xenium (blue, right) relative to snRNA-seq. Gene counts were averaged across WT samples and log_10_ -transformed counts per million (CPM). Density contours represent the distribution of genes, with increased color intensity indicating higher density. The diagonal line (slope = 1) denotes equality, and Pearson correlation coefficients (R) and the number of shared genes (n) between ST platforms and snRNA-seq are reported for each comparison. **G**. Computational workflow for *SpatialBench*, showing preprocessing, quality control, normalization, dimensionality reduction, and downstream analyses, with platforms color-coded across steps.

To account for resolution differences between ST platforms, we assessed data quality at binning resolutions of 8 µm and 16 µm and evaluated comparable bin numbers, transcript counts, and sparsity across resolutions (Fig. 1E). We benchmarked spatial gene profiles against both 10x Genomics FLEX and 3’ GEX references (Supplementary Fig. 1). Concordance was consistently higher with FLEX, which is probe-based and therefore more similar to Visium HD, while also showing closer comparability to targeted imaging platforms than 3’ GEX. Visium HD showed the strongest correlation (*R* = 0.92 across 14,999 shared genes), while MER-SCOPE (*R* = 0.73, *n* = 91) and Xenium (*R* = 0.64, *n* = 99) were evaluated on smaller targeted panels (Fig. 1F). FLEX was therefore used as the primary single-nuclei RNA seq (snRNA-seq) reference for benchmarking and downstream analysis. Our workflow was tailored to ensure fair comparisons, with all data processed in 8 µm and 16 µm bins for technical benchmarking, imaging-based datasets evaluated with multiple cell-level segmentation strategies, and downstream analyses applied uniformly to capture both technical performance and biological resolution (Fig. 1G).

### Detection sensitivity and background signal vary across ST platforms

To enable quantitative comparison across ST platforms, iST datasets were aggregated into 8 µm and 16 µm spatial bins and aligned to the Visium HD coordinate system using the matched H&E image as an anatomical reference (Fig. 2A, Supplementary Fig. 2). Spatial maps of transcript counts per bin showed consistent recovery of tissue architecture across platforms, whereas higher transcript densities in Visium HD reflected transcriptome-wide detection compared with targeted gene panels. Detection sensitivity was assessed using the 90 genes shared across platforms. Xenium showed the highest median transcript counts per bin (56 counts), followed by Visium HD (11 counts), whereas MERSCOPE showed substantially fewer counts (6 counts) (Fig. 2B). Detected genes per bin followed the same trend, with median values highest in Xenium (25 genes), intermediate in Visium HD (8 genes), and lowest in MERSCOPE (5 genes), consistent with higher detection sensitivity in Xenium and lower sensitivity in MERSCOPE.

**Fig. 2.**
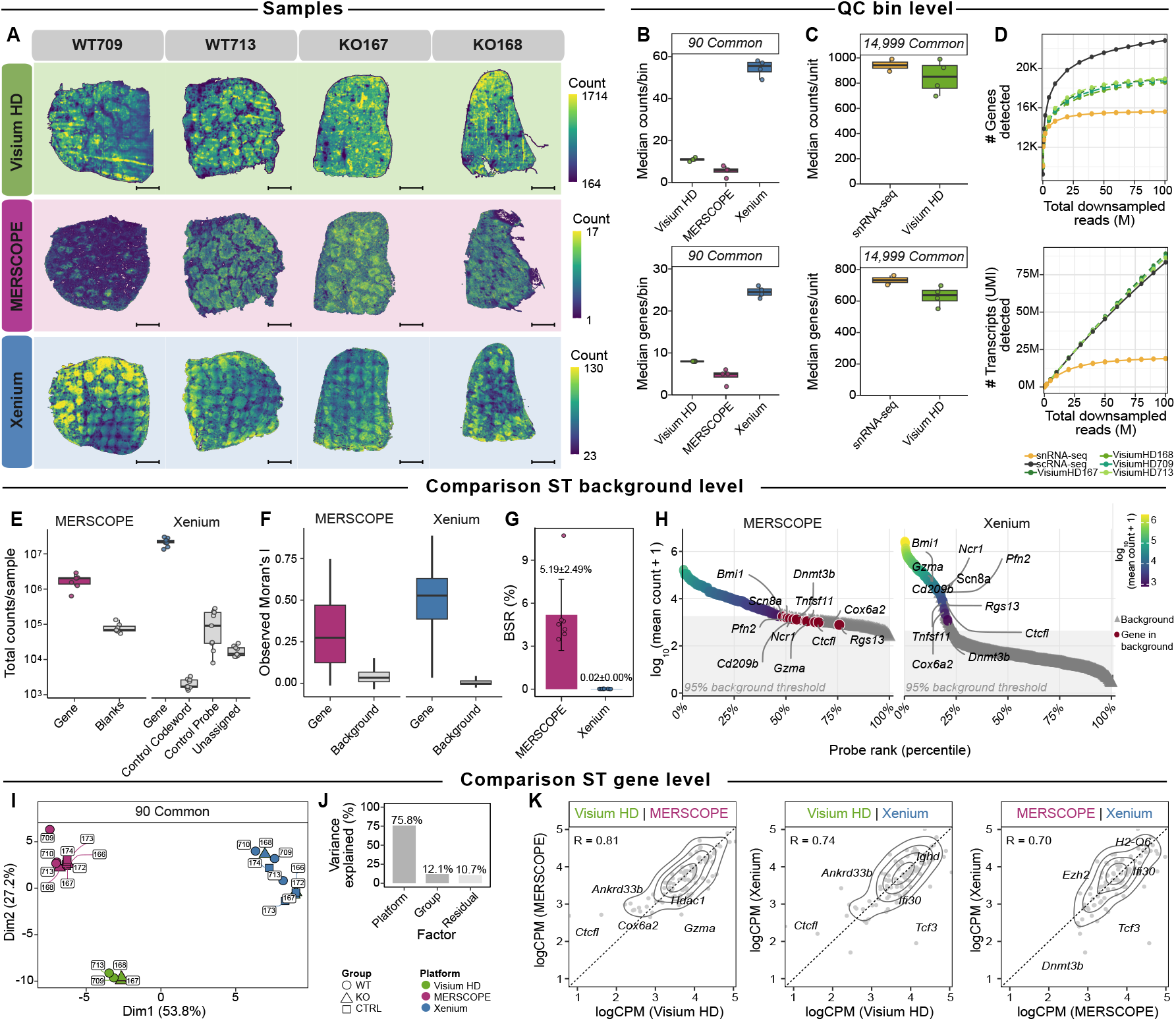
Comparative analysis of technical performance across harmonized high-resolution ST datasets. **A**. Matched spleen sections from wild-type (WT709, WT713) and conditional *Tbx21*knockout (KO167, KO168) mice profiled using Visium HD (green), MERSCOPE (magenta), and Xenium (blue). Heatmaps display total transcript counts per 8 µm bin, scaled per platform. Scale bars, 1 mm. **B**. Bin-level (8 µm) evaluation of median transcript counts per bin (top) and median detected genes per bin (bottom) for the 90 commonly shared genes across Visium HD, MERSCOPE, and Xenium plaforms. Points indicate individual samples. **C**. Bin-level (8 µm) evaluation of median transcript counts per bin (top) and median detected genes per bin (bottom) for the 14,999 commonly shared genes between Visium HD and snRNA-seq assays. Points indicate individual samples. **D**. Rarefaction curves showing the expected numbers of detected genes (top) and transcripts (Unique Molecular Identifiers; UMI) (bottom) in downsampled data as a function of mapped reads. M, millions. **E**. Total gene targeting and negative control probe counts per sample in MERSCOPE and Xenium. Box plots show sample-level totals (*n* = 9 per platform). Points indicate individual samples. **F**. Distributions of observed Moran’s I values for gene targeting probes and background probes (Blanks in MERSCOPE; Unassigned in Xenium) across all genes and samples, summarized by box plots. **G**. Background to signal ratio (BSR) of background probe detection across MERSCOPE and Xenium. Data are presented as mean *±* SEM. **H**. Rank-ordered mean probe counts (log_10_(mean count + 1)) for MERSCOPE and Xenium across all samples. Shaded area marks the platform-specific 95th percentile of background probe counts. Highlighted points indicate gene targeting probes at or below this threshold. **I**. Multidimensional scaling (MDS) of pseudo-bulk expression profiles for the 90 common genes across platforms. Point color denotes platform and shape denotes sample group. **J**. Variance partitioning of pseudo-bulk gene expression profiles attributed to platform (75.8%), biological group (12.1%), and residual (10.7%) variation. **K**. Cross-platform gene-level correlations. Density contour plots show pairwise comparisons of averaged log_10_CPM expression across shared genes in matched Visium HD, MERSCOPE, and Xenium samples, with Pearson correlation coefficients (R). Points indicate individual genes. Source data are provided as a Source Data file.

Compared with matched snRNA-seq data, Visium HD showed lower transcript recovery per bin and detected fewer genes, consistent with the reduced molecular capture efficiency typical of sST assays (Fig. 2C). To assess whether sequencing depth was sufficient across assays, rarefaction analysis of downsampled Visium HD, 3’ GEX (scRNA-seq) and FLEX (snRNA-seq) libraries revealed gene detection approaching saturation with increasing sequencing depth, while transcript counts continued to increase, consistent with expected library complexity and adequate sequencing depth (Fig. 2D).

We next quantified background signal and probe-level performance in iST platforms (Fig. 2E-H, Supplementary Fig. 3). Total counts per probe showed clear separation between gene-targeting probes and platform-specific negative control probes across all samples, with substantially higher counts observed for gene-targeting probes in both Xenium and MERSCOPE (median 21.8 M versus 1.7–91.3 K in Xenium; 2.0 M versus 69.1 K in MERSCOPE) (Fig. 2E), indicating strong separation between biological signal and technical background. Unassigned controls in Xenium and blanks in MERSCOPE, which lack complementary transcript targets, were used as an empirical estimate of assay background, as both are defined in the codebook without an associated target gene and therefore reflect spurious decoding events. Using these controls, spatial autocorrelation analysis using Moran’s I showed higher values for gene-targeting probes than for background probes in both platforms (Fig. 2F), indicating greater spatial structure of gene-targeting signal relative to assay background. Estimated background to signal ratio derived from background probes remained low, with mean values of 0.02% in Xenium and 5.19% in MER-SCOPE (Fig. 2G). Rank-ordered probe abundance distributions further showed that gene-targeting probes were largely separated from background levels, although MERSCOPE exhibited elevated background signal with a small subset of overlapping the assay background (Fig. 2H, Supplementary Fig. 4). These observations indicate clear separation between gene-targeting signal and assay background overall, while also revealing probe-specific variability in detection sensitivity, particularly among low-expressing probes in MER-SCOPE.

To enable robust gene-level comparison across platforms, pseudo-bulk expression profiles were generated by aggregating counts across spatial bins using the 90 genes shared between technologies. Multidimensional scaling showed that samples clustered primarily by platform, with platform identity accounting for 75.8% of total expression variance, whereas biological group accounted for a smaller proportion (12.1%) (Fig. 2I,J, Supplementary Fig. 5). Gene expression measurements were nevertheless positively correlated across technologies, with Pearson correlation coefficients ranging from *R* = 0.70 to *R* = 0.81 across pairwise comparisons (Fig. 2K), indicating substantial concordance in gene-level measurements despite platform-specific differences.

### Segmentation performance varies across methods in densely cellular spleen tissue

Accurate cell segmentation is critical to correctly assign transcripts to individual cells. The mouse spleen poses challenges to segmentation, due to its exceptionally high cellular density and tight spatial arrangement of transcriptionally distinct immune populations. We systematically evaluated segmentation performance across MERSCOPE (*n* = 9) and multi-modal Xenium (*n* = 5) datasets using platform-specific vendor default pipelines, Proseg^(36)^, a transcript-based refinement method, and Cellpose^(37,38)^, a deep learning–based segmentation method (Fig. 3). Vendor default segmentation differed between platforms and acquisition modes. MER-SCOPE used a pretrained Cellpose model on its proprietary cell boundary staining images, whereas Xenium unimodal datasets used nuclear segmentation followed by mask expansion, and Xenium multimodal datasets incorporated cell segmentation staining to define cell boundaries. To ensure comparability across segmentation strategies, analyses were restricted to multimodal Xenium datasets with segmentation staining, and unimodal vendor segmentation was applied to the same datasets to assess the impact of nuclear expansion. Representative tissue regions showed clear differences in cell boundary definition across segmentation strategies (Fig. 3A,C). Proseg generated enlarged masks that frequently encompassed adjacent cellular regions, consistent with transcript-driven reassignment within the initial vendor-defined segmentation, whereas vendor default segmentation produced smaller masks with variable boundary placement and often missed cell boundaries. Cellpose generated smaller, regularly shaped masks that more closely aligned with cellular morphology. These differences were quantified using cell count, cell area and transcript assignment metrics (Fig. 3B,D; Supplementary Fig. 6). In MERSCOPE datasets, Cellpose identified the greatest number of cells (total 402,026 per sample) compared to Proseg (270,145 cells) and vendor default segmentation (262,098 cells), consistent with smaller median cell areas (39.5 µm^2^ versus 49 and 53 µm^2^, respectively). Proseg yielded the highest transcripts per cell (median 36 versus 32 and 25 counts) and the highest transcript assignment rates (95% versus 79.3% and 86.6%). Comparable trends were observed in Xenium datasets. Cellpose again identified more cells (median 326,143 cells) than vendor default segmentation (166,161 cells) and Proseg (161,297 cells), whereas Proseg produced the highest transcripts per cell (376 versus 320 and 243 counts, respectively) and transcript assignment rates (99.3% versus 79.1% and 98.4%), demonstrating consistent effects of segmentation strategy on cell detection and transcript allocation across platforms.

**Fig. 3.**
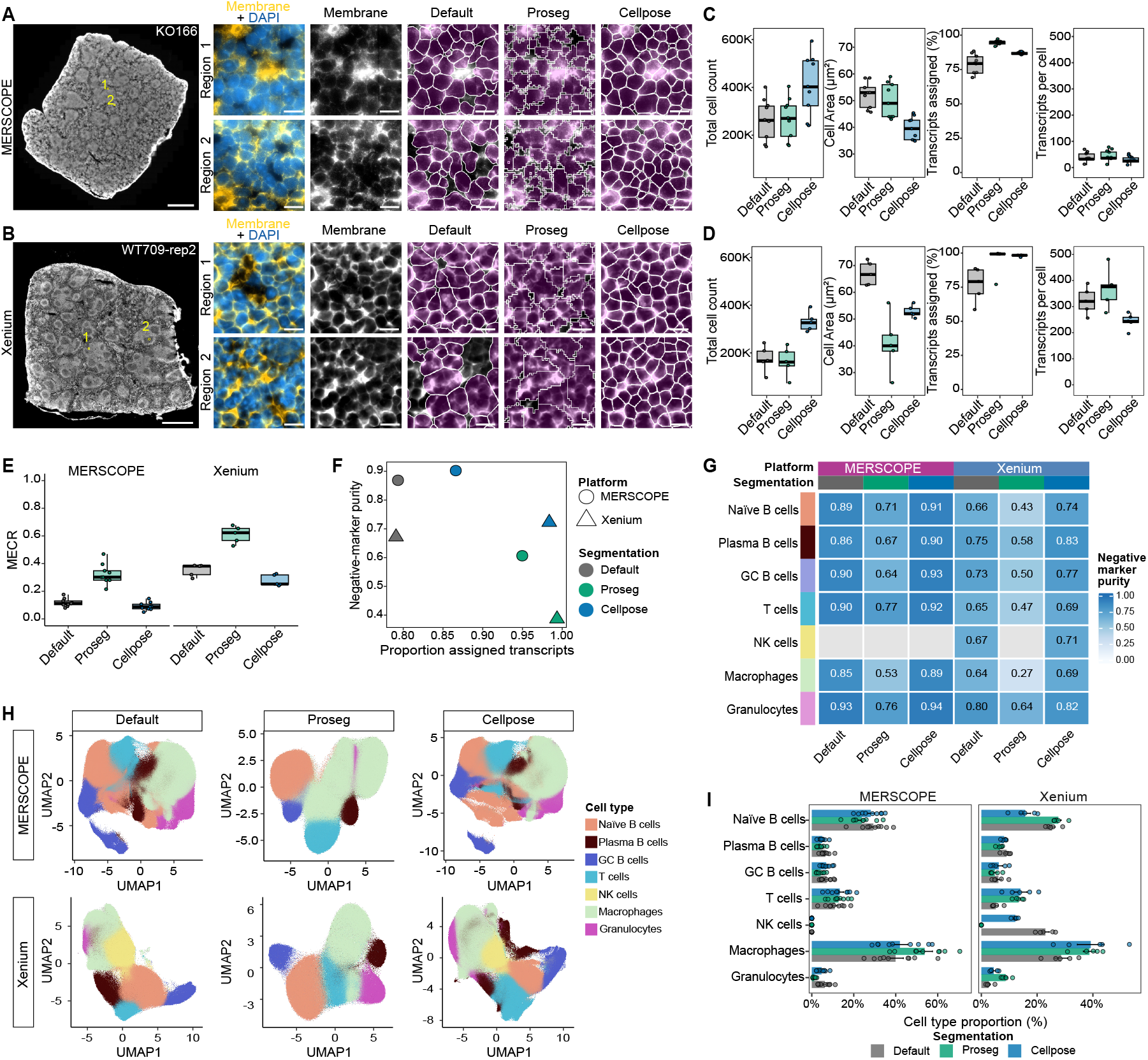
Comparison of cell segmentation methods across iST platforms. **A, B**. Representative (**A**) MERSCOPE tissue section (KO166) and (**B**) Xenium tissue section (CTRL174 rep2) with two regions of interest (ROIs) enlarged to visualise cell segmentation masks. Composite intensity images of cell membrane (yellow) and DAPI (blue) staining are shown alongside the membrane channel alone (grayscale). Segmentation outputs from the vendor default, Proseg, and custom Cellpose methods are displayed (magenta filled mask with white outlines) on top of membrane-only signal. Scale bar, 1 mm (overview) and 10 µm (zoomed ROI). **C, D**. Box plots of sample-level total cell count, median cell area, fraction of transcripts assigned to cells, and median transcripts per cell for(**C**) MERSCOPE (*n* = 9) and (**D**) Xenium multimodal (*n* = 5) samples for each segmentation method. Each point indicate individual sample. **E**. Box plots of sample-level mutually exclusive correlation rate (MECR) for each segmentation method in MERSCOPE and Xenium. Each point indicate individual sample. **F**. Relationship between the fraction of transcripts assigned to cells and negative marker purity across segmentation methods in MERSCOPE and Xenium. Each point represents a platform–segmentation combination, with platform indicated by shape and segmentation method by color. **G**. Heatmap of cell type specific negative marker purity across segmentation methods in MERSCOPE and Xenium. Exact values are displayed. **H**. Uniform Manifold Approximation and Projection (UMAP) of segmented single-cell expression profiles from MERSCOPE (top) and Xenium (bottom) for the vendor default (left), Proseg (middle), and custom Cellpose (right) segmentation methods, colored by annotated cell type. **I**. Bar plot of mean cells per sample by cell type and segmentation method in MERSCOPE and Xenium. Each point indicate individual sample. Data are presented as mean *±* SEM. Source data are provided as a Source Data file.

Segmentation fidelity metrics revealed consistent differences in transcript assignment accuracy across MERSCOPE and Xenium spleen datasets (Fig. 3E,F). Proseg produced the highest mutually exclusive co-expression rates (MECR; MERSCOPE 0.332; Xenium 0.676), consistent with admixing of mutually exclusive transcripts due to expanded cell boundaries. Vendor default segmentation showed intermediate MECR values (MERSCOPE 0.132; Xenium 0.406), whereas Cellpose consistently yielded the lowest MECR (MERSCOPE 0.106; Xenium 0.271), indicating improved separation of transcriptionally distinct splenic cell populations. Negative marker purity showed consistent ordering across both platforms, with Cellpose yielding the highest purity (0.902 in MERSCOPE and 0.723 in Xenium), followed by vendor default segmentation (0.869 and 0.672) and Proseg (0.606 and 0.387) (Fig. 3G,H). In line with this trend, UMAP embeddings revealed method-dependent organization of annotated cell types despite identical downstream processing (Fig. 3H). Segmentation strategy also influenced inferred cell-type abundances, with systematic shifts in cell-type proportions across methods (Fig. 3I). Together, these findings demonstrate that segmentation strategy strongly influences transcript assignment accuracy and downstream biological interpretation in densely cellular spleen tissue. Segmentation methods showed clear trade-offs, with Proseg maximizing transcript assignment at the expense of marker specificity, whereas Cellpose provided a better balance between transcript recovery and preservation of cell-type identity.

### Platform-dependent detection of genotype-associated *Tbx21* downregulation and transcriptional programs

To extend our comparison beyond technical performance to biological discovery potential, we leveraged the well-characterized phenotype of KO samples as a defined biological reference. In this model, Tbx21 loss in *Cd23*-expressing naïve and GC B cells disrupts GC organization by impairing their transition to a dark zone transcriptional program Ly et al. ^(39)^. Using matched Visium HD, MER-SCOPE and Xenium spleen sections from WT and KO mice, we compared cell-type annotation, spatial organization and genotype-associated transcriptional differences across platforms (Fig. 4).

**Fig. 4.**
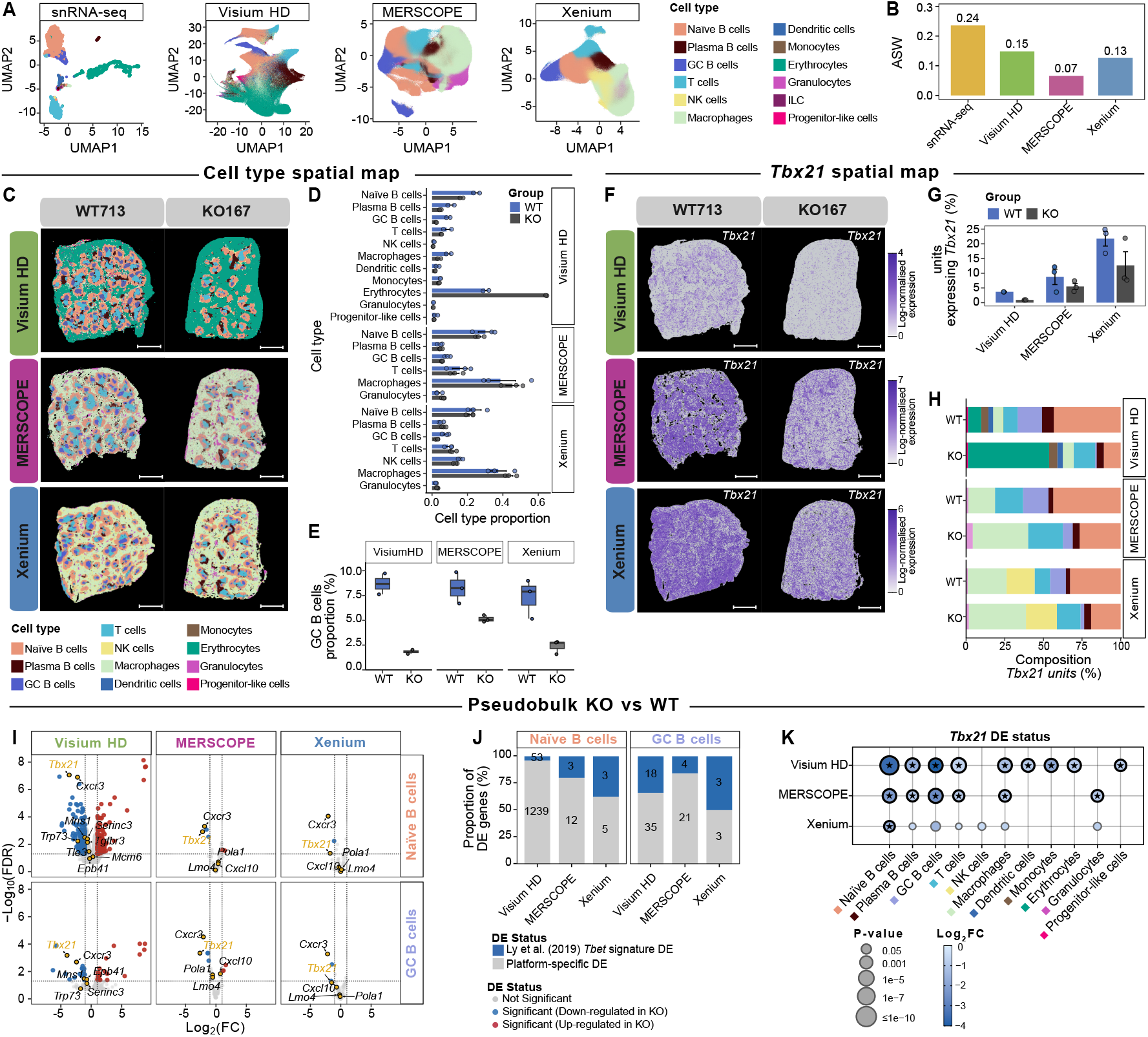
Cross-platform evaluation of ST analyses using a conditional *Tbx21* knockout ground truth. **A**. UMAP of snRNA-seq, Visium HD (8 µm bins), and cell-level MERSCOPE and Xenium data showing major spleen cell type annotations used for downstream analyses. **B**. Bar plot of average silhouette width (ASW) per platform, computed from the Principal component analysis (PCA) embeddings underlying the UMAP shown in Fig. 4A. **C**. Spatial maps of annotated cell types for WT713 and KO167 spleen sections across matched Visium HD (top), MERSCOPE (middle), and Xenium (bottom) samples. Each color represent a cell type. Scale bars, 1 mm. **D**. Bar plot of cell type proportions in WT and KO samples by platform. Visium HD includes WT and KO samples (*n* = 2 each), and MERSCOPE and Xenium include WT and KO samples (*n* = 3 each) (top). Data are presented as mean *±* SEM. **E**. Box plot of extracted GC B cell proportions from 4D in WT and KO samples by platform. Each point indicate individual sample. **F**. Spatial maps of *Tbx21* expression in WT713 and KO167 samples measured by Visium HD, MERSCOPE, and Xenium, with expression scaled within each platform. Scale bars, 1 mm. **G**. Bar plot of the percentage of *Tbx21*-positive units in WT and KO samples across platforms, defined as 8 µm bins for Visium HD and cells for MERSCOPE and Xenium. Data are presented as mean *±* SEM. **H**. Bar plot showing the cell type composition of *Tbx21*-positive units across platforms, colored by annotated cell type as in panel 4C. **I**. Volcano plots of pseudo-bulk KO versus WT differential expression in näive B cells (top) and germinal center B cells (bottom) across Visium HD, MERSCOPE, and Xenium. Genes are colored by differential expression status (|log_2_ FC| *>* 1, adjusted *P <* 0.05). *Tbx21* is highlighted in yellow with other key expected down-regulated genes. **J**. Bar plot of the proportion of significantly differentially expressed genes (adjusted *P <* 0.05) overlapping an external *Tbx21* (encodes for T-bet) knockout gene signature from Ly et al. ^(39)^ in näive B and germinal center B cells across platforms. **K**. Dot plot of pseudo-bulk *Tbx21* differential expression across annotated cell types for Visium HD, MERSCOPE, and Xenium. Dot color show log_2_ fold change (KO versus WT), with adjusted *P* -values indicated by dot size and annotated significance indicated by star symbol.

Consistent recovery of major immune populations across platforms provided the basis for these comparisons. Spatial datasets were analysed using BANKSY-based dimension reduction followed by UMAP embedding and spatial clustering (see Methods). For Visium HD, cell type labels were assigned using Robust Cell Type Decomposition (RCTD)^(40)^ and validated using canonical marker genes (Fig. 4A, Supplementary Fig. 7), while Xenium and MERSCOPE datasets were annotated using marker gene expression (Supplementary Fig. 8). Across all platforms, major splenic immune populations, including naïve, GC and plasma B cells, T cells, macrophages, dendritic cells and granulocytes, were consistently identified, enabling direct comparison of downstream analyses. Erythrocytes were detected only in Visium HD datasets because erythrocyte genes (*Hba/Hbb, Car1/Car2*, and *Klf1*) were not included in the targeted imaging-based panels owing to optical saturation constraints.

Transcriptional separation of these annotated populations varied across platforms but remained sufficient to support biological comparisons. Average silhouette width (ASW) values were lower for all spatial platforms than the matched snRNA-seq reference (0.24), with Visium HD (0.15) and Xenium (0.13) showing greater separation of immune populations than MERSCOPE (0.07) (Fig. 4B). This trend was consistent with spatial maps showing clearer immune compartment organization in Visium HD and Xenium compared to MERSCOPE (Fig. 4C). Despite differences in transcriptional separation, all platforms recovered broadly comparable immune compositions across WT and KO samples (Fig. 4D). Minor genotype-associated shifts were observed, but no platform showed major distortion of immune composition, indicating that downstream genotype comparisons were unlikely to be driven by platform-specific compositional biases. Notably, a consistent Tbx21-dependent reduction in GC B cell proportions was observed across platforms (Fig. 4E).

Having established comparable cell identity recovery and immune composition, we next examined whether the *Tbx21*^fl*/*fl^ *Cd23*-Cre perturbation was reflected in spatial gene expression (Fig. 4F, Supplementary Fig. 9). Across platforms, *Tbx21* signal was detected throughout tissue, with Visium HD showing more spatially structured and lower-intensity expression patterns. Quantification of *Tbx21*-positive units showed consistently higher proportions in WT than KO samples (Fig. 4G). In WT samples, *Tbx21* signal predominantly originated from B cell populations, whereas KO samples showed fewer positive units and broader distribution across annotated cell types (Fig. 4H), likely reflecting reduced signal specificity at low expression levels. Together, these results indicate that all platforms recover the expected direction of the biological perturbation.

We next tested whether these spatial differences were reflected in genotype-associated transcriptional changes. Pseudo-bulk differential expression analysis, aggregating expression profiles by annotated cell type and comparing KO and WT samples using limma–voom^(41,42)^, showed significant *Tbx21* downregulation in naïve B cells across all three platforms, whereas significant downregulation in GC B cells was detected only by Visium HD and MERSCOPE (Fig. 4I-K, Supplementary Fig. 10–11). In naïve B cells, *Tbx21* down-regulation was strongest in Visium HD (log_2_FC = −3.6, adjusted *P* = 8.6 × 10^−8^), followed by MERSCOPE (log_2_FC = −2.2, adjusted *P* = 1.0 × 10^−3^) and Xenium (log_2_FC = −1.8, adjusted *P* = 4.5 × 10^−2^). In GC B cells, significant downregulation was observed in Visium HD (log_2_FC = −4.0, adjusted *P* = 6.6 × 10^−4^) and MERSCOPE (log_2_FC = −2.6, adjusted *P* = 4.6 × 10^−4^), whereas Xenium showed a smaller, non-significant effect (log_2_FC = −1.5, adjusted *P* = 6.4 × 10^−2^). Beyond B cell compartments, significant *Tbx21* downregulation was detected in Visium HD and MER-SCOPE but not in Xenium, where significance was largely restricted to naïve B cells (Fig. 4I,K).

Finally, to determine whether these gene-level effects reflected broader T-bet-dependent transcriptional programs, we compared platform-specific pseudo-bulk results with an external *Tbx21* knockout gene signature from Ly et al. ^(39)^ (Fig. 4J). Visium HD identified a large number of differentially expressed genes in addition to *Tbx21*, including 1,239 platform-specific genes, although only a small fraction overlapped the reference signature. Imaging-based platforms identified fewer differentially expressed genes overall but showed higher proportional overlap with the reference signature. In naïve B cells, 20% of MERSCOPE and 37.5% of Xenium differentially expressed genes overlapped the external signature, whereas in GC B cells overlap was 16% and 50%, respectively. ROAST analysis further supported enrichment of the broader T-bet-dependent transcriptional program in Visium HD (ROAST *P* = 8.1 × 10^−4^ naïve B cells; *P* = 0.001 GC B cells) (Supplementary Fig. 12, Supplemental Table. 5), whereas equivalent analysis was not performed for imaging platforms due to gene panel limitations. Together, these analyses show that while *Tbx21* downregulation is consistently detected across spatial platforms, the breadth of associated transcriptional programs detected depends on platform gene coverage.

### Germinal center dark and light zone organization is differentially captured across spatial platforms

To evaluate the ability of high-resolution ST technologies to resolve GC architecture, we identified GC regions across Visium HD, MERSCOPE and Xenium datasets in WT and T-bet conditional KO mouse spleens (Fig. 5A, Supplementary Fig. 7–8). GC regions were defined based on spatial localization and expression of canonical GC markers (*Aicda, Bcl6, Cd83, Cxcr4* and *Rgs13*). Consistent with known GC organization, *Bcl6* showed broad GC expression, *Aicda* was enriched in dark zone (DZ) regions, and *Cd83* preferentially localized to light zone (LZ) regions (Fig. 5A)^(43,44)^. To resolve DZ and LZ compartments, GC B cells were subclustered using spatially informed BANKSY embeddings and curated zonal markers. In Visium HD, DZ clusters were enriched for proliferation and DNA repair genes (including *Rrm1, Cdca8, Ccna2, Myc* and *Aicda*), whereas LZ clusters showed enrichment of activation and antigen presentation markers (including *Cd38, Cd83, Cd86, Ciita* and *Irf4*; Supplementary Fig. 13). Xenium data similarly resolved DZ and LZ states despite reduced gene coverage (Fig. 5A, Supplementary Fig. 14). In contrast, MERSCOPE did not show reproducible zonal segregation, consistent with its lower transcript detection sensitivity (Fig. 2A).

**Fig. 5.**
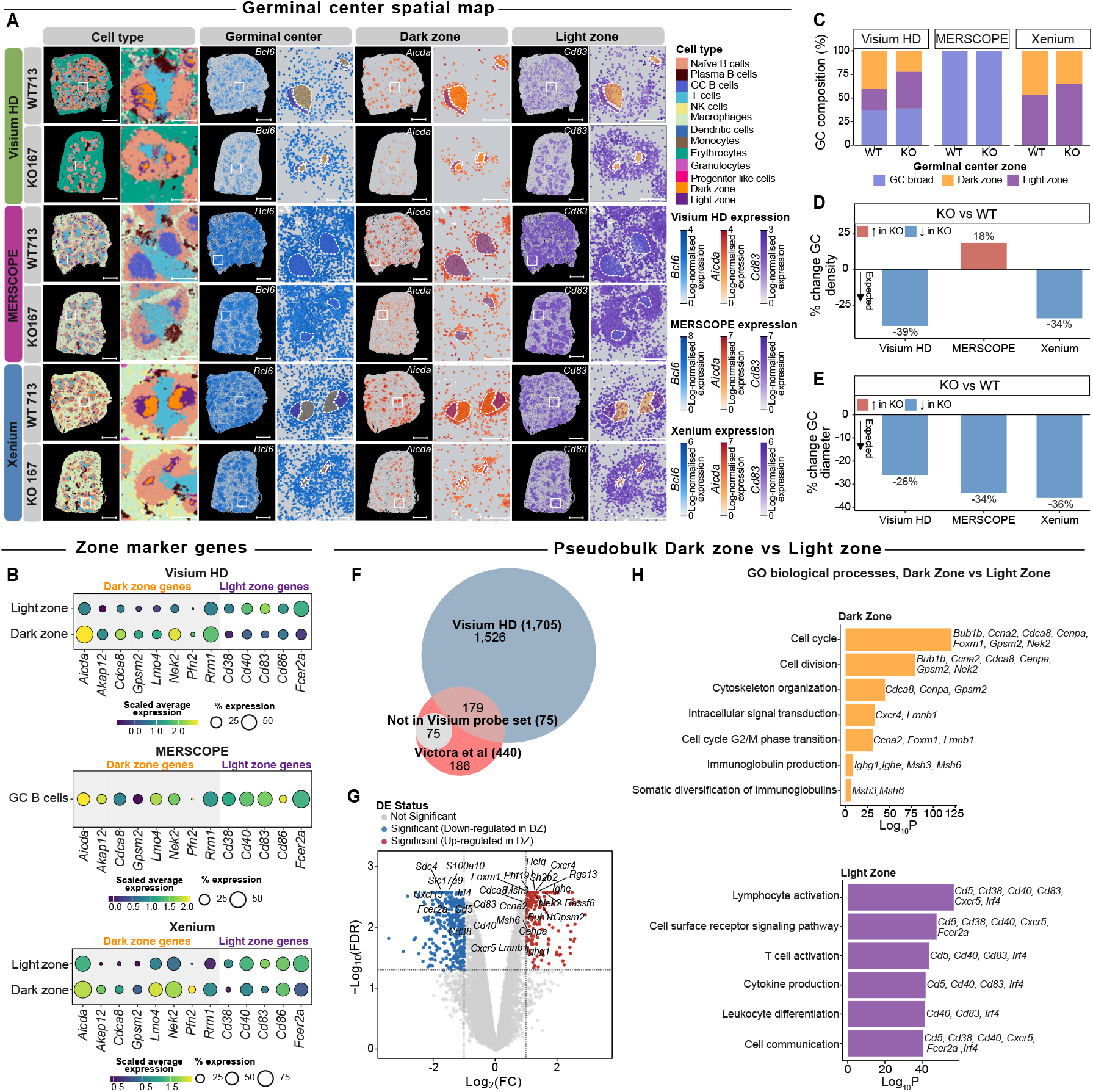
Interrogating germinal center spatial organization with high resolution ST platforms. **A**. Spatial maps of annotated cell types for WT713 and KO167 spleen samples profiled by Visium HD, MERSCOPE, and Xenium. Germinal centers (GC) are delineated with dark and light zones annotated, and representative marker gene expression is shown for *Bcl6* (GC), *Aicda* (dark zone), and *Cd83* (light zone). Dotted outlines indicate GC boundaries. Color intensity indicates log-normalised expression. Scale bars, 1 mm (overview) and 200 µm (ROIs). **B**. Dot plots of dark zone and light zone marker gene expression across platforms. Average expression is shown by color and the percentage of expressing units is indicated by dot size. Visium HD and Xenium show expression stratified by annotated dark and light zones, while MERSCOPE shows expression within GC B cells. **C**. Bar plot of GC composition in WT and KO samples for Visium HD, MERSCOPE, and Xenium, showing the relative contribution of dark zone, light zone, and unassigned GC regions. Bars represent mean proportions across samples, with colors indicating GC sub-compartments. **D-E**. Percentage change in (**D**) GC density and (**E**) GC diameter in KO relative to WT samples across platforms. **F**. Venn diagram showing the overlap of differentially expressed genes (adjusted *P* < 0.05) between dark and light zones in WT samples and a published GC zone gene set from Victora et al. ^(43)^. **G**. Volcano plot of pseudo-bulk differential expression between dark and light zones in WT samples profiled with Visium HD. Genes are colored by differential expression status using | log_2_ FC| > 1 and adjusted *P* < 0.05. **H**.Gene Ontology (GO) biological process enrichment for genes differentially expressed between dark and light zones in WT samples. Bar plots show significantly enriched GO terms for dark zone (top) and light zone (bottom) gene sets, ranked by enrichment significance (log_10_*P*). Representative contributing genes are shown for each term. Source data are provided as a Source Data file.

Cross-platform comparison using shared DZ and LZ markers showed consistent relative enrichment patterns in Visium HD and Xenium (Fig. 5B). Consistent with the previously described transcriptional continuum between DZ and LZ states, in which a substantial proportion of GC B cells occupy intermediate positions at any given time^(45,46)^, zonal differences were primarily reflected as relative expression shifts rather than compartment-exclusive markers, with greater transcriptional mixing observed in LZ regions. Spatial organization was also anatomically consistent, with DZ regions positioned adjacent to T cell zones and LZ regions located closer to naïve B cell follicles (Fig. 5A). Together, these results indicate robust GC zonal resolution in Visium HD and consistent recovery in Xenium, whereas MERSCOPE did not support reliable zonal separation. GC composition differed between WT and T-bet conditional KO samples across platforms (Fig. 5C). In Visium HD, GC B cells were classified as DZ, LZ or unassigned GC, likely reflecting this continuum of intermediate transcriptional states and potentially compounded by mixed states captured within 8*µ*m bins (Supplementary Fig. 13A). KO samples showed a relative increase in LZ states and a reduction in DZ states compared with WT. Xenium data showed a similar shift toward increased LZ representation without an intermediate category, reflecting its single-cell resolution. MERSCOPE analyses were therefore restricted to total GC B cells due to lack of zonal resolution.

To quantify GC organization independently of zonal assignment, we performed region-level GC segmentation and morphological analysis (Fig. 5D,E, Supplementary Fig. 15). WT samples showed greater total GC area than KO samples in Visium HD and Xenium, whereas trends were less consistent in MERSCOPE (Fig. 5D). GC morphology analysis further showed broader GC diameter distributions in WT samples, including larger structures exceeding 200,µm, whereas KO samples showed more restricted size distributions (Fig. 5E, Supplementary Fig. 15C). Together, these results indicate that loss of T-bet is associated with coordinated changes in GC zonal composition and tissue-scale GC organization, with consistent trends across Visium HD and Xenium.

To characterize transcriptional differences between dark and light zone compartments, we aggregated Visium HD expression profiles for annotated DZ and LZ GC B cells in WT mice and performed pseudo-bulk differential expression analysis using the limma–voom^(41,42)^ framework (Fig. 5F,G, Supplemental Table. 6). Robust zonal annotation combined with the broad gene coverage of Visium HD enabled well-powered differential expression testing, whereas analogous analyses were not performed for Xenium due to the targeted nature of the gene panel. We identified differentially expressed 829 DZ genes, and 876 LZ genes in WT mice. This was 1526 number greater genes as compared to 440 genes identified by Victora et al. ^(43)^ (Fig. 5F, Supplementary Fig. 16). Gene set enrichment using the gene ontology (GO) biological processes database as a reference set and the 829 DZ and 876 LZ genes as test sets were performed in limma^(41)^ (Fig. 5H). As expected, the most significant GO terms enriched in DZ genes represented numerous processes involved in proliferation and cell cycle progression (Fig. 5H, top). Within these pathways, several DZ-defining genes with various roles in proliferation, cell cycle, and mitosis were present (*Bub1b, Ccna2, Cdca8, Cenpa, Gpsm2, Nek2*)^(43)^. Of note, the transcriptional activator *Foxm1*, a master regulator of cell proliferation^(47)^, was present across several terms enriched in GC B cells (Fig. 5H, top). Moreover, the intracellular signal transduction GO term, which was enriched in this gene set, contained the gene *Cxcr4* that drives GC DZ polarization^(44)^, and *Lmnb1* that encodes important epigenetic regulator of DZ B cell somatic hypermutation^(48)^. Despite lower significance, GO terms associated with immunoglobulin production (p = 4.49 × 10^−9^) and somatic diversification of immunoglobulins (p = 1.36 × 10^−6^) were enriched in GC DZ genes (Fig. 5H, top). Genes encoding the constant region of the immunoglobulin heavy chain (*Ighg1, Ighe*) and genes encoding mismatch repair proteins (*Msh3, Msh6*) were present within these pathways. The most significant GO terms enriched in LZ genes encompassed key processes occurring in the GC LZ such as cell differentiation and activation, cell sur-face receptor signalling, cytokine production, and cell communication (Fig. 5H, bottom). Several LZ defining genes, including *Cd38, Cd83*, and *Fcer2a*^(43)^ were present within these terms. Moreover, the gene *Cd40*, which promotes positive selection of high affinity antigen receptors on B cells^(49)^, the genes *Ccr6* and *Irf4*, which indicate GC LZ B cell differentiation into memory B cell precursors^(50,51)^, and the chemokine *Cxcr5* that indicates recruitment of B cells to the GC LZ, was present across several of these GO terms enriched for GC LZ genes in WT mice (Fig. 5H, bottom). Thus, while dark and light zone structure can be resolved across platforms, transcriptome wide profiling with Visium HD is required to interrogate germinal center zonal transcriptional programs in an unbiased manner.

## Discussion

Here we present a biologically controlled benchmark of three high-resolution ST platforms (Visium HD, Xenium and MERSCOPE) using matched malaria-challenged spleen tissue to determine how platform design influences recovery of biological structure at near-cellular resolution. While recent ST benchmarking studies have compared platform performance across tissues^(25,29,30,33)^, many rely on unmatched samples or focus primarily on technical performance metrics rather than biological interpretability. By integrating matched spatial datasets with FLEX snRNA-seq, and 3’ GEX scRNA-seq from our prior *SpatialBenchVisium* resource^(35)^, we established a unified immune reference in which platform differences could be evaluated against a defined *Tbx21*-driven biological perturbation rather than technical benchmarks alone. This design enables assessment of how platform characteristics influence the recovery and interpretation of known immune biology, providing a more practical basis for evaluating ST performance.

Across platforms, differences in transcript detection, gene recovery and background signal were evident, but these did not prevent recovery of major immune structure or core *Tbx21*-associated biological effects. *Tbx21* encodes the transcription factor T-bet, a well-established regulator of GC B cell differentiation and zonation, and its conditional deletion provides a reproducible perturbation of GC organization and transcriptional programs^(39)^. Recovery of these expected T-bet–dependent effects across platforms therefore provides a biologically grounded benchmark of ST performance. Platform differences instead primarily influenced the resolution at which these biological programs could be interpreted rather than whether they could be detected. This suggests that commonly used QC metrics may overestimate the biological impact of technical differences and reinforces that benchmarking within defined biological systems provides a more meaningful assessment of ST performance than technical comparisons alone.

Consistent with previous benchmarking efforts describing trade-offs between gene coverage and spatial resolution^(30,33)^, our results highlight complementary strengths between sequencing- and imaging-based platforms. Visium HD enabled transcriptome-scale characterization of GC transcriptional programs and pathway-level analysis, whereas imaging platforms provided improved resolution of immune organization at cellular scale. Rather than identifying a single optimal platform, these findings emphasize that ST technologies differ primarily in the scale at which biological questions can be addressed. Resolving fine-grained immune structures such as GC dark and light zones remains a recognized challenge for ST technologies. Kleshchevnikov et al. ^(52)^ showed that DZ and LZ B cells are among the most difficult populations to accurately map using Bayesian deconvolution of standard resolution Visium data from human lymph node, and our previous *SpatialBenchVisium* study identified GC regions in the same malaria-challenged spleen model but lacked sufficient resolution to resolve zonal organization^(35)^. Here, our *SpatialBench* resource, incorporating near-cellular resolution platforms, enables direct spatial analysis of GC zonation without reliance on deconvolution.

The germinal center provided a stringent biological test of platform capability. The dense cellular architecture and subtle transcriptional gradients between GC zones place simultaneous demands on transcript detection sensitivity, gene coverage and accurate transcript assignment. Under these conditions, Visium HD and Xenium consistently resolved GC zonation whereas MERSCOPE showed reduced recovery of these distinctions. Given the similar gene panels used for iST platforms, this difference likely reflects transcript detection efficiency rather than fundamental platform limitations. These comparisons demonstrate that accurate GC zonation analysis depends on both spatial resolution and transcript detection performance, indicating that spatial resolution alone is insufficient. More broadly, these results illustrate how platform characteristics translate into differences in biological resolution when analyzing complex immune tissue structure.

Our results further demonstrate how analytical choices can shape biological conclusions. In densely cellular spleen tissue, segmentation strategy substantially influenced transcript assignment and inferred lineage specificity. While the importance of segmentation is well recognized^(31,32)^, our data show that reliance on default vendor segmentation may be insufficient in immune tissues with closely apposed cell populations. Custom imaging-based segmentation reduced transcriptional mixing and improved lineage specificity, indicating that segmentation optimization is essential when interpreting cell-type–resolved analyses in such tissues. These findings align with recent work emphasizing the need to explicitly model transcript misassignment and contamination in spatial datasets^(53,54)^.

Generating matched multi-platform ST datasets proved technically demanding, requiring substantial experimental optimization, with several datasets not meeting quality thresholds (Supplemental Table 1). These practical constraints, together with the cost and experimental complexity of spatial assays, likely explain the limited availability of directly comparable multi-platform benchmarks. These factors also represent important limitations of the present study. Imaging analyses were performed using earlier Xenium and MERSCOPE gene panels available at the time of data generation, whereas newer expanded panels such as Xenium 5K and the MER-SCOPE 1K panel now provide substantially broader gene coverage that may improve biological resolution. In addition, MERSCOPE data were generated prior to Vizgen’s updated gene imaging chemistry (v2 chemistry), which was reported to substantially enhance transcript detection sensitivity^(55)^. Reliance on nuclear segmentation in some Xenium datasets may also underestimate improvements achievable through multimodal-staining segmentation approaches, particularly in densely cellular immune tissues. Future bench-marking efforts should also extend to whole-transcriptome in situ platforms, including Atera (10x Genomics)^(56)^ and CosMx (Bruker Spatial Biology)^(57)^, which overcome the gene panel constraints inherent to the imaging-based platforms evaluated here.

Despite these limitations, *SpatialBench* provides a biologically grounded reference dataset for evaluating high-resolution ST technologies in immune tissue. Beyond platform comparison, integration of matched spatial datasets with complementary single-cell and single-nucleus references within a well-characterized GC model enables investigation of B cell differentiation, GC zonation and *Tbx21*-dependent immune programs. The resource also supports future benchmarking work of spatial analysis methods, including segmentation, deconvolution and spatial domain detection, by providing a system with defined cellular composition and transcriptionally defined GC zonation, establishing an interpretable reference for assessing whether computational methods recover consistent biological structure across platforms. As ST technologies and computational methods continue to evolve, multi-platform resources anchored in well-defined biology such as *SpatialBench* will be essential for reproducible benchmarking and for understanding how platform design influences biological conclusions.

## Methods

### Sample preparation

#### Mouse spleen samples

Mice were housed at the animal facility of the Walter and Eliza Hall Institute of Medical Research (WEHI) in plastic cages under controlled environmental conditions (20–22^◦^C, 40–70% relative humidity) with a 14 h light / 10 h dark cycle. All animal experiments were performed in accordance with institutional and national ethical guidelines and were approved by the WEHI Animal Ethics Committee. Male *Tbx*21*fl/flCd*23*Cre* mice, in which *Tbx21* is deleted in mature follicular B cells, together with *Cd*23*Cre* control mice and 8-week-old wild-type (WT) female C57BL/6J mice, were infected intravenously with 1 × 10^5^ *Plasmodium berghei* ANKA–infected red blood cells to induce splenic germinal center formation. Following the development of clinical symptoms, mice were treated with chloroquine and pyrimethamine to clear infection, as described previously^(39)^.

#### Tissue preparation

Twelve days post-infection, mice were euthanised and spleens were either snap frozen in Optimal Cutting Temperature (OCT) compound using a PrestoCHILL instrument (Milestone), or fixed in 10% (v/v) neutral buffered formalin and processed into formalin-fixed paraffin-embedded (FFPE) blocks using standard histological procedures at the WEHI Advanced Histotechnology facility.

### Single-cell and single-nuclei RNA-seq data generation

#### 10X 3’GEX experiments

scRNA-seq data from mouse spleen generated from the same infection model were obtained from our previously published dataset (Du et al. ^(35)^; GEO accession: GSE254652) and used as matched single-cell references for ST benchmarking. Experimental details are described in Du et al. ^(35)^.

#### 10X FLEX experiments

Five 25 µm scrolls were taken from each FFPE spleen blocks and dissociated into single nuclei according to the 10x Genomics pestle dissociation demonstrated protocol (CG000632, Rev A). Following trituration, 1.5 million nuclei were immediately subjected to overnight probe hybridization. Samples were further processed according to the NextGem Chromium Fixed RNA Profiling Single-plexed Samples User Guide (CG000691, Rev A). For each sample, a target of 10,000 nuclei was captured.

### Spatial transcriptomic data generation

#### Gene panel design and construction

MERSCOPE (Vizgen) and Xenium (10x Genomics) technologies require the use of a predefined gene panel. Based on prior single-cell and spatial data analysis^(35)^ and expert knowledge, we selected genes useful for cell type identification and linked to malaria-associated inflammation biology. For MERSCOPE, we excluded genes that were too abundant (>1,000 FPKM) or too short (<750 nt in length) that were potentially challenging for MERFISH imaging. We also adjusted probeset number for highly abundant genes for Xenium imaging. The steps resulted in a 94-gene and 100-gene panel for MERSCOPE and Xenium, respectively. Gene panel contents are provided in Supplemental Table 2.

#### Xenium experiments

Xenium *in situ* expression analysis was conducted following the manufacturer’s instructions. All reagents, including water, were molecular-grade nuclease-free. Slide incubation steps were performed by positioning a Xenium Thermocycler Adaptor (10x Genomics, PN-3000954) on a thermal cycler (Bio-Rad C1000 Touch) to ensure efficient and even heat transfer. Sample preparation began by sectioning fresh-frozen tissues at 10 *µ*m thickness on a CryoStar NX70 cryostat (Epredia) and placing them on the Xenium slide (10x Genomics, PN-1000460). Slides were stored at -80^◦^C for ≤4 weeks until processing.

The Xenium slides were prepared following the Xenium In Situ fresh-frozen Tissue Demonstrated Protocol (CG000579, Rev F). In brief, slides were incubated at 37°C for 1 min, fixed with formaldehyde, permeabilised with methanol, and assembled into Xenium cassettes for the remainder of the preparation. Probes hybridization, ligation and amplification were then performed according to the 10x Genomics’ Demonstrated Protocol (CG000582, Rev H). To enhance cell segmentation accuracy, selected samples (CTRL172, CTRL173, CTRL174, WT709, WT710) were stained overnight with Xenium Cell Segmentation Add-on Kit (10x Genomics, 1000662) as described in the manufacturer’s protocol (CG000749, Rev B). This kit provides a cocktail of stains for broad-coverage cell segmentation across four imaging channels, targeting nuclei (DAPI), cell membranes (antibodies for ATP1A1, E-Cadherin, CD45), the cytoplasm (18S rRNA), and cytoskeletal proteins (alphaSMA/Vimentin). Slides stained with the add-on kit underwent additional steps of stain enhancement before treated with autofluorescence quenching and nuclei staining. Finalised slides were stored in 0.05% Tween-20/PBS in the dark at 4^◦^C for ≤7 days before loading onto the Xenium Analyser instrument.

The Xenium Analyser is a fully automated instrument for spatial imaging of gene and protein targets within tissue section at subcellular resolution. We followed the Xenium Analyser User Guide (G000584, Rev K) to prepare consumables and operate the instrument. Briefly, up to two Xenium slides and the required reagents were loaded onto the instrument, and the appropriate codebook matching the gene panel was selected. A low-resolution, whole-slide image was then generated for users to mark regions of interest for spatial profiling. Data were acquired in iterative cycles of probe hybridization with fluorescent labels, image capture, and probe stripping under automated sample and liquid handling. The Xenium Onboard Analysis pipeline was run directly on the instrument for image processing, cell segmentation, image registration, decoding, deduplication and secondary analysis. We acquired Xenium data on 15 samples in three separate batches. Batch 1 samples (WT713, CTRL173-rep1, KO166, KO167) were acquired using instrument software version 1.7.6.0 and Onboard Analysis pipeline version 1.7.1.0. Batch 2 samples (WT709-rep1, WT710-rep1, CTRL172, CTRL174-rep1, KO168) were acquired using instrument software version 1.8.2.1 and Onboard Analysis pipeline version 1.7.1.0. Batch 3 samples (WT709-rep2, WT710-rep2, CTRL172-rep2, CTRL173-rep2, CTRL174-rep2) were acquired using instrument software version 3.1.0.0 and Onboard Analysis pipeline version 3.1.0.4.

After the run, slides were removed from the Xenium Analyser instrument and treated with 10mM sodium hydrosulfite solution to remove quenching chemicals according to 10x Genomics’ Demonstrated Protocol (CG000613, Rev B). This was followed by Mayer’s Hematoxylin and Eosin Y staining, dehydration and coverslipping. Histology images were taken on the SLIDEVIEWTM VS200 research slide scanner (Olympus Life Science).

#### MERSCOPE experiments

Fresh-frozen mouse spleens were cut at 10 µm thickness onto MERSCOPE slides (Vizgen, 1050001) using the Feather S35 microtome blade (ProSciTech, UDM-S35) and CryoStar NX70 cryostat (Epredia). Slides were kept inside the cryostat for 15 min and then transferred to a hot plate set at 37^◦^C for 1 min to enhance adherence of the section to the glass slide. Samples were then fixed in freshly prepared 4% PFA/PBS and permeabilised by 70% ethanol at 4^◦^C overnight. Slides were then placed in a Photobleacher instrument (Vizgen, 1010003) for autofluorescence quenching for at least 4 h at room temperature. Slides were stored for up to 4 weeks at 4^◦^C before proceeding to the next step. To visualise and segment individual cell boundaries, sections were stained with the cell boundary kit (Vizgen, 10400118) before being subjected to encoding probe hybridization for 36 h in a 37^◦^C humidified incubator. After hybridization, the samples were embedded in a thin polyacrylamide gel for RNA anchoring, then treated with a clearing mix supplemented with proteinase K (NEB, P8107S, 1/100 dilution) for 2–4 h at 37^◦^C until no visible tissue was evident in the gel.

The MERFISH imaging process was performed according to the MERSCOPE Instrument User Guide (Vizgen, number 91600001). In brief, an imaging kit was thawed in a 37^◦^C water bath for 60 min, activated, and loaded into the MER-SCOPE instrument. The slide was stained with DAPI and PolyT Staining Reagent, assembled into the flow chamber, and connected to the fluidic lines in the MERSCOPE instrument. The fluidics were then activated to fill the flow chamber with liquid without air bubbles. A low-resolution image for the DAPI and PolyT stains was acquired at 10 × magnification, and regions of interest (ROIs) were manually drawn in the MERSCOPE software. The instrument then switched to a 60 × /1.4 NA oil immersion objective for automated image acquisition and fluidic control for iterative rounds of readout probe hybridization and removal. Images were collected at eight 1.5 µm-thick *z*-stacks over a total thickness of 12 µm. Readout probes were imaged in the 560 nm, 650 nm, and 750 nm channels, and DAPI and PolyT stains were imaged in the 405 nm and 488 nm channels, respectively. To image 94 genes and cell boundary stains, each run consisted of six rounds of imaging for an 18-bit MERFISH experiment, followed by three sequential rounds of three-color FISH imaging. The total MERFISH imaging time was approximately 24 h for each experiment.

#### Visium HD experiments

Fresh-frozen mouse spleens were cut at 10 *µ*m thickness onto SuperFrost Plus slides (Epredia) using a Feather S35 microtome blade (ProSciTech, UDM-S35) and a CryoStar NX70 cryostat (Epredia). SuperFrost Plus slides were pre-cooled in the cryostat prior to sectioning and kept within the cryostat until all sectioning was complete. Sections were taken from four different mouse spleens (WT709, WT713, KO167, and KO168), and after sectioning, slides were stored in sealed slide mailers at −80 ◦C for ≤4 weeks until processing. The sections were fixed, followed by H&E staining and imaging according to the Visium HD fresh-frozen Tissue Prep Handbook (CG000763, Rev A) by following the H&E Staining & Imaging workflow. Visium libraries were then prepared according to the Visium HD Gene Expression User Guide (CG000685, Rev B). All slide incubation steps were performed by positioning a Visium Low Profile Thermocycler Adapter (10x Genomics, PN-3000823) on a thermal cycler (Bio-Rad C1000 Touch) to ensure efficient and even heat transfer. Libraries were sequenced on the NextSeq 2000 (Illumina) according to the 10x Genomics sequencing guidelines.

### Single-nuclei RNA-seq data analysis

Data from 10x FLEX experiments were automatically demultiplexed to their sample of origin. Empty droplets were identified and removed using *emptyDrops* in DropletUtils^(58,59)^ (v1.28.1). Cells with fewer than 100 total transcripts were removed to exclude low-quality captures. Quality control outliers were identified using a three median absolute deviations (MADs) approach (*perCellQCFilters*; scater^(60)^, v1.36.0), removing cells with low numbers of detected genes or total transcripts, or high mitochondrial content. Discarded cells were inspected to confirm that no biologically relevant populations were removed. Expression values for retained cells were normalized using *logNormCounts* in scran^(61)^ (v1.36.0). The top 10% of highly variable genes (*getTopHVGs*; scran) were used for principal component analysis (PCA), and the first 10 principal components were used for louvain clustering. Cell types were annotated using SingleR^(62)^ (v2.10.0) in per-cell mode with the matched 3’ GEX scRNA-seq dataset as reference, followed by manual refinement based on canonical marker gene expression.

### Spatial transcriptomics data analysis

#### Generation of bin-level spatial expression profiles

Bin-level spatial expression profiles were generated for all datasets to enable direct comparison across platforms. Visium HD gene-by-bin count matrices at 8 µm resolution were generated using 10x Genomics SpaceRanger (v3.1.2) and used to construct Seurat objects (Seurat v5.1.0) for downstream analysis. For iST platforms (Xenium and MERSCOPE), bin-level Seurat objects were constructed directly from decoded transcript coordinates using custom R functions. Xenium transcripts were read from *transcripts*.*parquet* and filtered by molecule quality score (QV ≥ 20). MERSCOPE transcripts were read from *detected_transcripts*.*csv* in micron coordinate space. Transcripts were assigned to 8 µm and 16 µm spatial bins based on their spatial coordinates and aggregated per bin and gene to generate gene-by-bin count matrices. Platform-specific negative control features, comprising negative control probes, negative control codewords, and unassigned codewords for Xenium, and blanks for MERSCOPE, were processed using the same binning procedure and stored as separate assays to enable background signal quantification. Bin centroid coordinates were derived from bin indices and represented as square polygons using the Seurat functions *CreateCentroids* and *CreateFOV*, and associated with the corresponding bin-level Seurat objects.

#### Spatial alignment across platforms

Imaging-based bin centroids were aligned to the Visium HD coordinate system for matched tissue sections (WT709, WT713, KO167, and KO168) using STalign^(63)^ (v1.0; Python), accessed from R via reticulate (v1.42.0). Bin-level expression profiles were rasterized using *STalign*.*rasterize* (dx = 30) and aligned to the matched Visium HD H&E image following intensity normalization with *STalign*.*normalize*. An initial affine transformation was estimated from manually defined landmark correspondences using *STalign*.*L_T_from_points* and *STalign*.*to_A*, and subsequently refined using diffeomorphic large deformation diffeomorphic metric mapping (LDDMM) via *STalign*.*LDDMM* (500 iterations; *σ*_*P*_ = 0.2, *σ*_*M*_ = 0.2, *σ*_*B*_ = 0.3, *σ*_*A*_ = 0.3; *diffeo_start*=100). The final transformation was applied to bin centroid coordinates using *STalign*.*transform_points_source_to_target*.

#### Cell segmentation and generation of cell-level expression profiles

Cell boundaries and transcript-to-cell assignments were obtained using vendor-provided segmentation pipelines, independently trained image-based segmentation with Cellpose (v2.3.2) ^(37,38)^, and transcript-based segmentation with Proseg (v2.0.4)^(36)^.

Vendor-provided segmentations were used as the reference segmentation for each platform. For MERSCOPE datasets, segmentations and transcript-to-cell assignments were obtained using the Vizgen Post-Processing Tool (VPT v1.3.0) via the WEHI-SODA-Hub/spatialvpt pipeline (v1.0.0), which applies a Cellpose-based segmentation model to cell boundary (Cellbound), nuclear (DAPI), and cytoplasmic (Poly-T) staining channels. For Xenium datasets, segmentations and transcript-to-cell assignments were obtained using Xenium Ranger (v2.0.1), which defines cell boundaries using DAPI-based nuclear segmentation followed by isotropic nuclear expansion (5 µm). Segmentation stains were disabled so that all Xenium datasets, including multi-modal acquisitions, were segmented using an identical nuclear expansion procedure.

Image-based segmentation was performed using Cellpose. For MERSCOPE datasets, a Cellpose model was fine-tuned from the pretrained cyto2 model using manually annotated masks from 39 representative fields of view. Training images were generated from nuclear (DAPI) and cell boundary (Cellbound2 and Cellbound3) staining channels at imaging plane *z* = 3. Ground-truth masks were generated by manual refinement of pretrained cyto2 segmentations using the Cellpose interface, and images were contrast-normalized using contrast-limited adaptive histogram equalization prior to training and inference. The trained model was applied to all MERSCOPE datasets to generate cell boundary polygons. For Xenium datasets, the Cellpose model was retrained using 11 manually annotated Xenium image regions, together with down sampled MERSCOPE training data to match Xenium pixel resolution. Segmentation was performed using QuPath (v0.5.1) with the BIOP Cellpose extension (v0.9.6)^(64,65)^. Predicted cell boundary polygons were imported into Xenium Ranger (v3.1) to regenerate transcript-to-cell assignments based on the updated segmentation. Training and inference parameters are provided in Supplemental Table 3.

Transcript-based segmentation was performed using Proseg, which refines cell boundaries based on transcript spatial distributions. Proseg was run with default settings for MER-SCOPE (*proseg –merscope*) and Xenium (*proseg –xenium*) datasets, using vendor-provided segmentations for initialization. Proseg refines existing cell boundaries without introducing new cells, and resulting segmentations represent refinement of the initial segmentation. Cell boundary polygons were generated using the *proseg-to-baysor* utility with default settings.

For all segmentation methods, transcript-to-cell assignments were generated by assigning decoded transcript coordinates to the corresponding cell boundary polygons, and cell-by-gene count matrices were constructed for downstream analysis.

#### Quality control and preprocessing

ST datasets were processed using a consistent computational framework, with platform-specific preprocessing where required to enable direct comparison across platforms. Cells or spatial bins outside tissue boundaries were excluded. For iST datasets, low-confidence segmentations were removed using a transcript count threshold defined as the 10th percentile of cells and exceeding the 90th percentile of background probe signal (*<* 6 detected transcripts per cell). For Visium HD datasets, a low-quality spatial region in WT709 identified by reduced transcript density was excluded (Supplementary Fig. 2A).

#### Normalization and feature selection

Expression values were normalized using platform-appropriate methods. Visium HD datasets were normalized using relative counts normalization (*normalization*.*method = “RC”, scale factor = 5,000*). Highly variable genes were identified independently for each Visium HD sample using Seurat’s *FindVariableFeatures* function, and the union of 3,836 genes was retained for dimensionality reduction and spatial clustering. All genes were retained for imaging-based datasets due to the targeted panel design.

#### Spatial feature construction and integration

Visium HD datasets were processed jointly using normalized expression values following the BANKSY recommendations. Briefly, BANKSY neighbourhood matrices were constructed from variable genes (*k*_geom_ = 6), followed by BANKSY PCA and UMAP using *λ* = 0.2 to optimise spatial resolution .

For iST datasets, spatially informed feature representations were constructed using BANKSY^(66)^ (v1.1.0). Expression values were scaled prior to feature construction, and BANKSY was applied independently to each sample to preserve within-section spatial structure. Resulting features were combined for joint analysis using Harmony^(67)^ (v1.2.3) with sample identity specified as the integration variable. BANKSY was run using *λ* = 0.3 and *k*_geom_ = 12.

#### Clustering and cell type annotation

For Visium HD datasets, we used the 8 µm gene-by-bin count matrices and assigned cell-type annotation using Robust Cell Type Decomposition (RCTD) in SpaceXr^(40)^ (v1.1.0) using doublet mode. The annotated matched wild-type snRNA-seq data were used as reference for RCTD. Annotation accuracy was supported by concordance between transferred labels, marker gene expression, and expected spatial organization (Supplementary Fig. 7).

For iST datasets, dimensionality reduction was performed using principal component analysis (PCA) on BANKSY-derived features, retaining the top 30 principal components.

A shared nearest neighbour (SNN) graph was constructed from Harmony-corrected embeddings using the first 20 principal components, and clusters were identified using the Leiden algorithm (resolution = 0.6). Two-dimensional embeddings were generated using uniform manifold approximation and projection (UMAP) applied to Harmony-corrected principal components. Clusters were annotated based on canonical marker gene expression and spatial localization (Supplementary Fig. 8).

#### Germinal center zonation analysis

For Visium HD datasets, to resolve dark and light zone spatial domains beyond annotations derived from the snRNA-seq reference, we performed BANKSY clustering on bins annotated as germinal center B cells, analysing WT and KO samples separately (resolution = 0.5). Spatial domains were defined based on expression of an expanded panel of established DZ and LZ marker genes (Supplementary Fig. 13), which recapitulated the expected germinal center spatial organisation (Fig. 5A).

Germinal center zonation was resolved in imaging-based datasets by targeted subclustering of GC B cells identified in the initial analysis. Clustering was repeated on the subset of GC B cells using BANKSY-derived features with Harmony integration and Leiden clustering (resolution = 0.2). Subclusters were annotated as dark zone (DZ) or light zone (LZ) based on established marker expression, including *Aicda, Akap12, Cdca8, Gpsm2, Lmo4, Nek2, Pfn2, Rrm1* for DZ and *Cd38, Cd40, Cd83*, and *Fcer2a* for LZ (Supplementary Fig. 14).

#### Pseudo-bulk differential expression analysis

Pseudo-bulk differential expression was performed separately for each cell type by aggregating counts at the biological replicate level. For Visium HD datasets, counts were aggregated across 8 µm spatial bins assigned to each cell type, with two biological replicates per genotype (*n* = 2). For imaging-based datasets, counts were aggregated at the cell level using custom Cellpose-based segmentation for MERSCOPE and vendor-provided segmentation for unimodal Xenium datasets, with three biological replicates per genotype (n = 3). Analyses were restricted to WT and KO samples, and cell types with fewer than two biological replicates per genotype were excluded.

Genes with insufficient expression were filtered using *filterByExpr* (edgeR v4.2.2)^(68)^, and library size and composition were normalized using the trimmed mean of M-values method (*calcNormFactors*; edgeR)^(69)^. Mean–variance relationships were modeled using *voom* (limma v3.60.6)^(41,42)^, and gene-wise linear models were fit using *lmFit* with genotype as the explanatory variable. Contrasts were specified using *contrasts*.*fit* to compare KO and WT samples, and statistical significance was assessed using empirical Bayes moderation with *eBayes*. Adjusted *P* values were computed using the Benjamini–Hochberg procedure, and genes with adjusted *P <* 0.05 were considered differentially expressed.

An additional pseudo-bulk differential expression analysis was performed for Visium HD datasets to compare DZ and LZ regions within WT samples by aggregating counts across 8 µm bins assigned to each zone and applying the same limma–voom framework described above.

#### Pathway enrichment analysis

Gene set testing was performed on pseudo-bulk KO–WT differential expression results using ROAST^(70)^ implemented in limma. A germinal center zonation signature comprising 109 genes from Ly et al. ^(39)^ was tested for coordinated differential expression within each cell type. ROAST was applied within the same limma–voom linear modeling framework and KO–WT contrast used for differential expression analysis, using rotation-based testing with Benjamini–Hochberg correction to control the false discovery rate; gene sets with adjusted *P <* 0.05 were considered significant.

Gene ontology (GO) enrichment analysis was performed for WT Visium HD DZ–LZ differential expression results using over-representation analysis implemented in limma. Enrichment of GO biological process terms was assessed using *goana* (*species = “Mm”*), which applies a hypergeometric test relative to the tested gene universe, and terms were ranked using *topGO*. Statistical significance was determined using Benjamini–Hochberg correction, with adjusted *P <* 0.05 considered significant. Enrichment was performed separately for genes upregulated in DZ and LZ as defined by WT pseudo-bulk DZ–LZ differential expression.

### Benchmarking metrics

#### Binned data quality control

Bin-level quality control metrics were computed from gene-by-bin count matrices at 8 µm and 16 µm resolution. Let *x*_*gb*_ denote the observed count for gene (or probe) *g* in spatial bin *b*. For cross-platform benchmarking, analyses were restricted to the 90 genes shared across platforms; platform-specific analyses additionally used the full gene panel or transcriptome where applicable.

#### Bin-level transcript abundance and gene detection

For each spatial bin *b*, total transcript counts was defined as

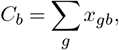

that is, the total number of detected transcripts in that bin.

Gene detection per bin was defined as

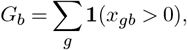

representing the number of unique genes with at least one detected transcript in bin *b*.

*C*_*b*_ therefore quantifies local transcript density, whereas *G*_*b*_ reflects gene complexity within each spatial bin.

#### Matrix sparsity

Matrix sparsity was defined as the proportion of zero entries in the gene-by-bin matrix,

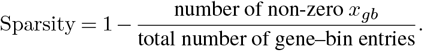

Sparsity provides a global measure of matrix density and signal dispersion.

#### Background signal and probe-level performance

Background signal was quantified using platform-specific negative control features. For Xenium datasets, background comprised unassigned transcripts, control codewords, and control probes. For MERSCOPE datasets, background comprised blank probes only. All remaining decoded probes were considered target genes.

### Total target and background burden

Total target and background counts per sample were defined as

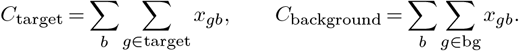

These totals were computed per sample and compared across platforms on a log_10_ scale.

### Probe-level signal relative to background

For each probe *g*, total counts were summed across spatial bins within each sample. The platform-level mean signal per probe was then calculated as the average total count across samples,

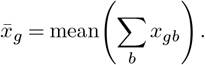

An empirical background threshold (*T*_bg_) was defined as the 95th percentile of mean signal among background probes,

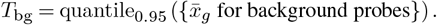

Target probes with 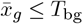 were classified as overlapping the background distribution.

### Spatial autocorrelation of target and background probes

Spatial structure was quantified using Moran’s *I*. For probe *g*,

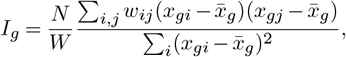

where *x*_*gi*_ denotes expression of probe *g* in spatial unit *i, w*_*ij*_ are spatial weights between locations *i* and *j, N* is the number of spatial units, and *W* = ∑_*i,j*_ *w*_*ij*_.

Moran’s *I* was computed on rasterized bin-level expression profiles using MERINGUE (v1.0)^(71)^. Spatial neighbors were defined using *getSpatialNeighbors* (*filterDist* = 100). Statistical significance was assessed using *filterSpatialPatterns* with Benjamini–Hochberg correction (*adjustPv = TRUE, α* = 0.05), and probes were required to be detected in at least 5% of spatial units (*minPercentCells* = 0.05). Distributions of Moran’s *I*_*g*_ were compared between target and background probes.

### Background-to-signal ratio

Background contamination in imaging-based platforms was quantified using negative control features. Total background calls (*C*_bg_) were defined as the sum of counts across all negative control features. The number of negative control features was denoted *N*_bg_. Total target gene calls (*C*_target_) were defined as the sum of counts across all target genes, and the number of target genes was denoted *N*_target_.

The background-to-signal ratio (BSR) was defined as

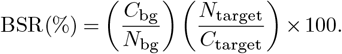

BSR was computed per sample and summarised per platform.

#### Cross-platform gene expression concordance

Cross-platform concordance was assessed using Pearson correlation of per-gene mean expression profiles. Raw pseudo-bulk counts were TMM-normalised (edgeR), converted to counts per million (CPM), and log_10_(CPM+1)-transformed. For each platform, gene-level expression was averaged across WT samples, and Pearson correlation coefficients were computed between platform pairs across the shared gene set.

### Segmentation performance evaluation

#### Mutually exclusive correlation rate (MECR)

MECR was computed to quantify marker mixing between distinct cell types. For marker genes assigned to different cell types, co-detection was defined as the proportion of cells in which both markers were detected among cells in which at least one marker was detected,

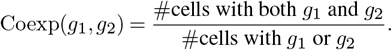

MECR was defined as the mean co-detection rate across marker pairs representing distinct cell types, following SpatialQM package^(72)^.

#### Negative marker purity

Negative marker purity was calculated as described by Marco Salas et al. ^(32)^. Gene–cell type pairs with reference positive fraction *<* 0.005 were classified as negative markers. The purity score was defined as

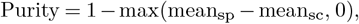

where mean_sp_ and mean_sc_ denote mean positive fractions across negative marker pairs in spatial and single-cell data, respectively.

#### Concordance of germinal center spatial organization

GC architecture was reconstructed from annotated spatial coordinates for each platform. Spatial bin or cell centroids annotated as germinal center populations were converted to binary masks to define GC regions within the tissue. Morphological filtering and connected-component labeling were performed using EBImage (v4.46.0)^(73)^ to identify individual GC structures per sample (Supplementary Fig. 15). For each GC, morphological features were computed including area (µm^2^), equivalent diameter. GC count and density (GCs per mm^2^) were calculated per sample relative to total tissue area.

## Data and Code Availability

Spatial and single nuclei data are available in the BioStudies database (http://www.ebi.ac.uk/biostudies) under accession number S-BSST2361. The *SpatialBench* data details are available at https://github.com/ashsolano/SpatialBench. The code for the analysis can be found at https://github.com/ashsolano/SpatialBench. The code for running VPT segmentation for MERSCOPE can be found at https://github.com/WEHI-SODA-Hub/spatialvpt.

## Ethics approval

All experiments were performed with ethics approval from the Animal Ethics Committee of the Walter and Eliza Hall Institute of Medical Research.

## Consent for publication

Not applicable.

## Author Contributions

A.S. and C.W. planned and performed data analysis, generated figures, interpreted results and wrote the manuscript. R.K.H.Y. planned and generated data, advised on data analysis and interpretation and wrote the manuscript. D.A.-Z. planned and supervised data collection and analysis, interpreted results and wrote the manuscript. C.J.A.A., L.L. and R.M. generated data. P.R., I.Z., A.M., M.C., Y.X., Y.P., S.S., P.F.H., L.W., C.J.S. and Y.C. contributed to study design, performed data analysis and interpreted results. L.J.I. provided tissue samples and contributed to study design. K.G-J. provided mouse models and interpreted results. H.W.K. advised on data analysis and interpreted results. K.L.R., D.S.H., R.B and M.E.R. designed the study, supervised data generation, analysis and interpretation and wrote the manuscript. All authors read and approved the final manuscript.

## Acknowledgements

We thank the WEHI Advanced Genomics Facility, Center for Dynamic Imaging and Advanced Histotechnology facility for supporting the generation and analysis of single-cell and spatial transcriptomics data for this project.

