## Supplementary Information for "*SpatialBench*: Comparative cross-platform benchmarking of high-resolution spatial transcriptomics using matched mouse lymphoid tissue"

### Supplementary Figures and Information from “*SpatialBench*: Comparative cross-platform benchmarking of high-resolution spatial transcriptomics using matched mouse lymphoid tissue”

#### Supplementary Figures

8 Cell-type marker gene expression across annotated populations in MERSCOPE and Xenium . 10
9 Spatial expression of Fcer2a (Cd23) across spatial transcriptomics platforms . . . 11
10 Cell-type resolved differential expression of *Tbx21* in Visium HD knockout and wild-type samples . . . 12
11 Cell-type resolved differential expression of *Tbx21* in imaging-based spatial transcriptomics datasets . . . 13
12 *Tbx21* (Tbet) signature enrichment across cell types in Visium HD samples . . . 14
13 Expression of germinal center dark and light zone marker genes and spatial zonation in Visium HD samples. . . . 15
14 Identification and spatial mapping of germinal center dark and light zones in Xenium samples. 16
15 Germinal center identification and morphological comparison across spatial transcriptomics platforms . . . 17
16 Dark and light zone gene signature enrichment in Visium HD samples . . . 18

### List of Tables

1 *SpatialBench* sample metadata and processing information. . . . . . . . . . . . . . . . . . . . 19
2 Gene panel composition for MERSCOPE and Xenium assays. . . . . . . . . . . . . . . . . . . . 19
3 Cellpose segmentation parameters for imaging-based spatial transcriptomics data. . . . . . . . 20
4 Differential expression results for Visium HD knockout versus wild-type across annotated cell types. . . . . . . . . . . . . . . . . . . . . . . . . . . . . . . . . . . . . . . . . . 21
5 ROAST gene set concordance with Ly et al.[1] using Visium HD KO versus WT differential expression results. . . . . . . . . . . . . . . . . . . . . . . . . . . . . . . . . . . . . . 21
6 Differential expression results between germinal center dark and light zones in Visium HD wild-type samples. . . . . . . . . . . . . . . . . . . . . . . . . . . . . . . . . . . . . . 22

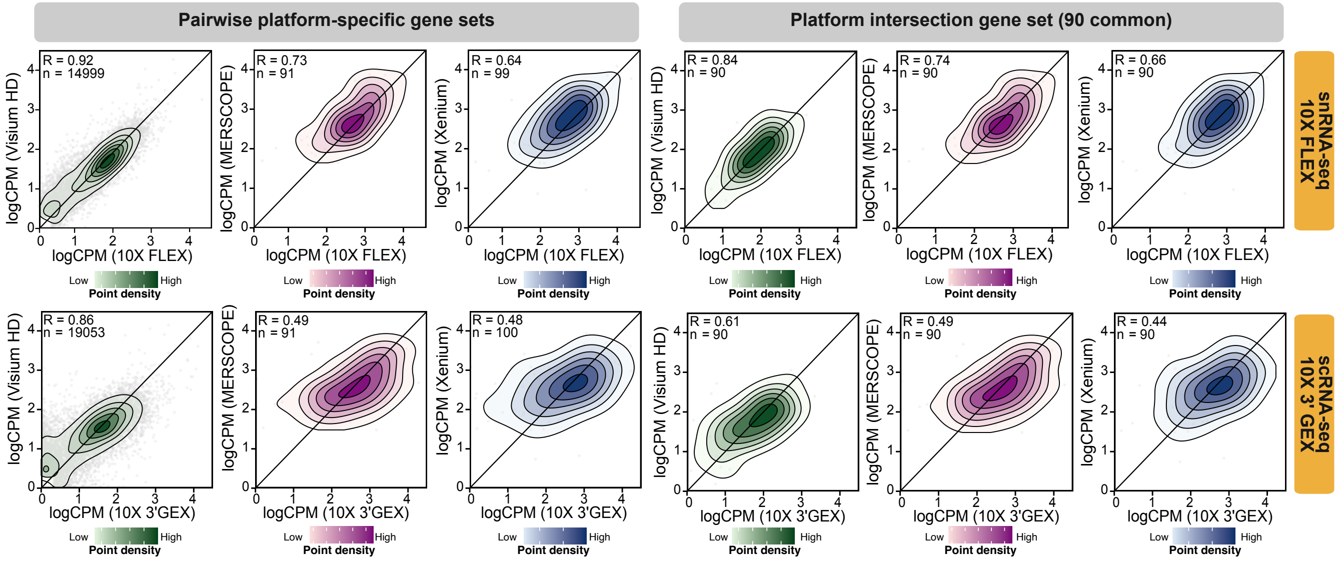

**Supplementary Fig. 1: Correlation of spatial transcriptomics platforms with snRNA-seq FLEX and scRNA-seq 3' GEX references. A.** Pearson correlation of gene expression profiles between Visium HD, MERSCOPE and Xenium spatial datasets and reference transcriptomic profiles generated using 10x Genomics snRNA-seq FLEX (top) or scRNA-seq 3' GEX (bottom). Correlations were calculated across all detected genes (left) and the shared 90-gene panel (right). Each panel reports the Pearson correlation coefficient ($R$) and the number of detected genes ($n$). Density contours represent the distribution of genes, with increased color intensity indicating higher density.

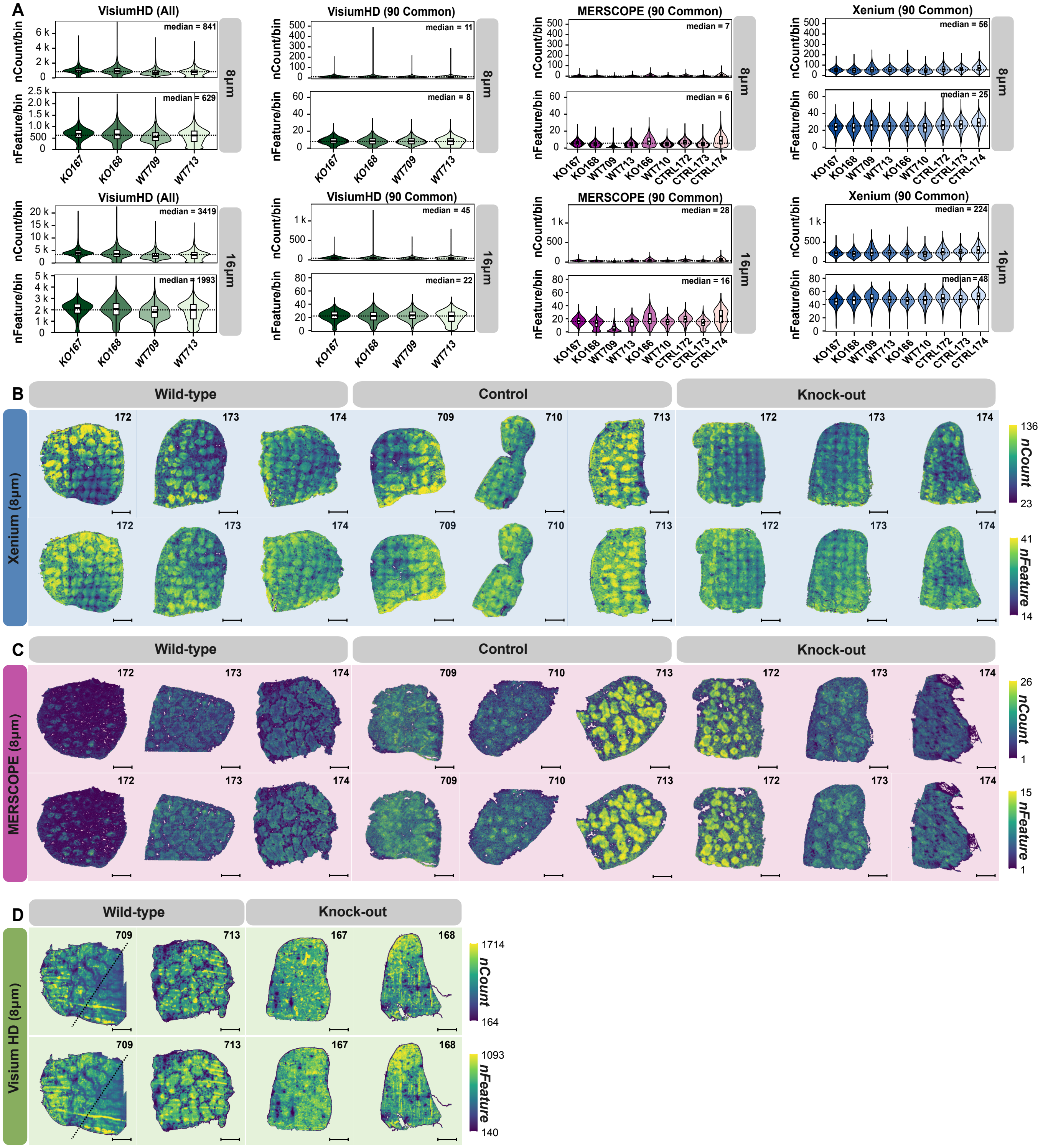

**Supplementary Fig.** 2: **Transcript and gene count distributions across spatial transcriptomics platforms at 8 µm and 16 µm bin resolution. A.** Per-bin distributions of total transcripts (*nCount*, top row) and detected genes (*nFeature*, bottom row) for Visium HD, MERSCOPE and Xenium. Results are shown for two spatial aggregation scales (8 µm and 16 µm bins). For Visium HD, values are reported for all detected genes (left) and for the cross-platform shared 90-gene panel (right). **B-D**. Spatial maps of Xenium (**B**), MERSCOPE (**C**) and Visium HD (**D**) datasets (8 µm bins) showing transcript counts (*nCount*, top row) and detected genes (*nFeature*, bottom row), arranged by genotype (wild-type, control and knockout) with sample identifiers above each map. Colour scales indicate the dynamic range used within each platform. Scale bars, 1 mm.

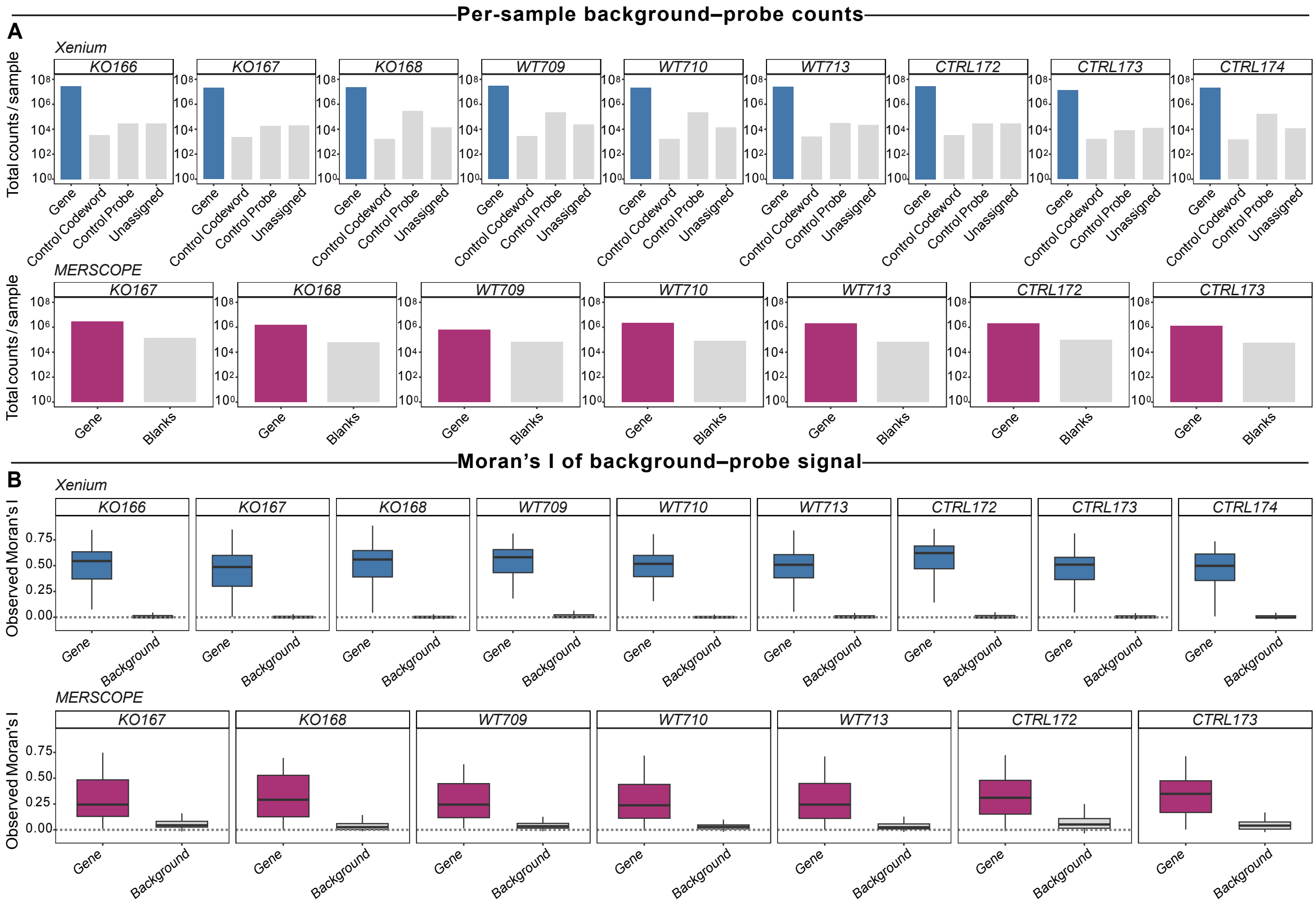

**Supplementary Fig.** 3: **Negative control signal and spatial autocorrelation in imaging-based spatial transcriptomics platforms. A.** Per-sample negative control signal. Total counts assigned to manufacturer-designated negative controls for Xenium (top row) and MERSCOPE (bottom row). **B.** Spatial autocorrelation of negative control signal measured using Moran's I. Boxplots show the distribution of observed Moran's I values across negative controls for Xenium (top row) and MERSCOPE (bottom row).

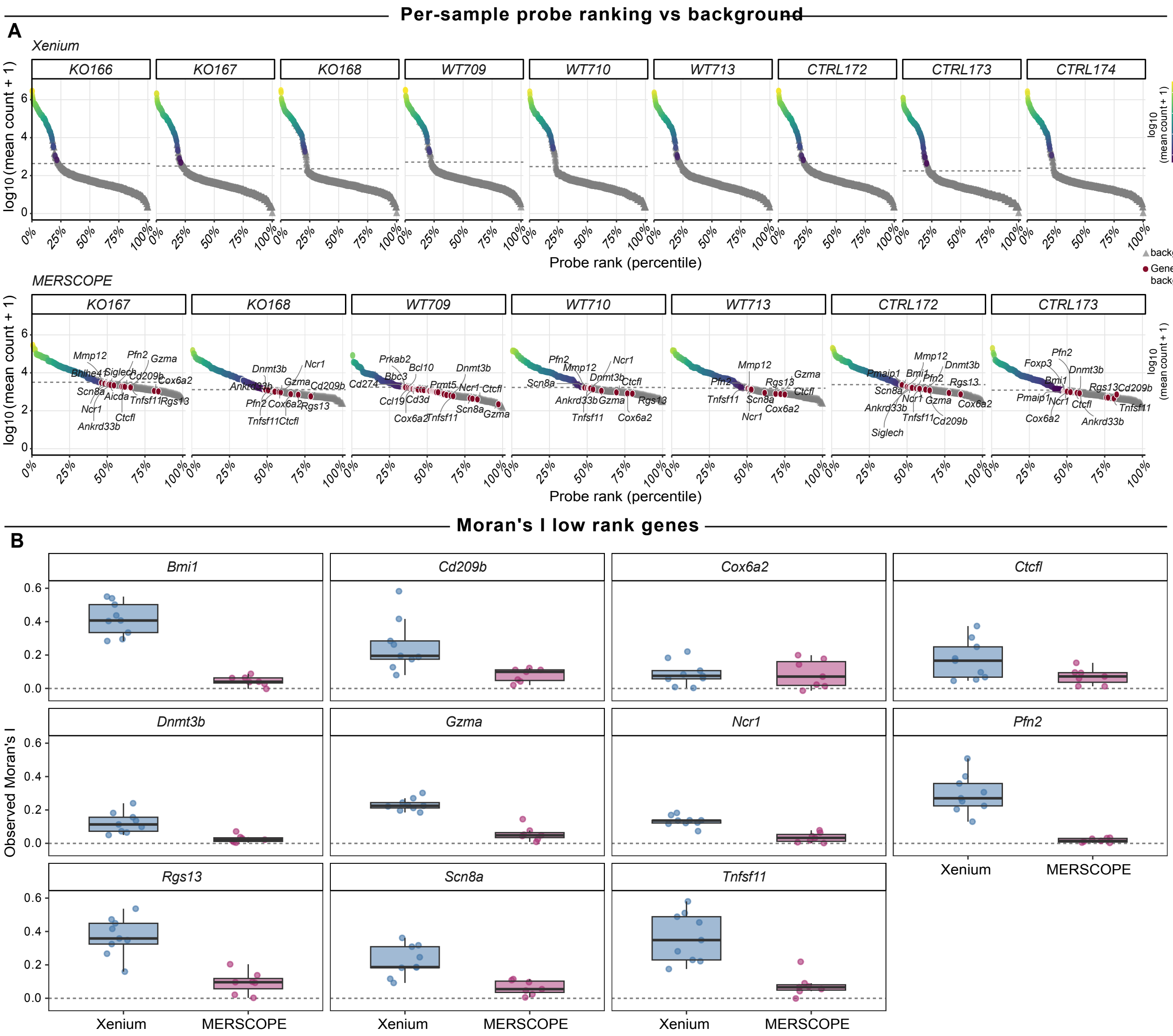

**Supplementary Fig.** 4: **Probe signal distributions relative to background controls and spatial autocorrelation in Xenium and MERSCOPE. A.** Rank plots of probe signal for each sample, shown for Xenium (top row) and MERSCOPE (bottom row). Each point represents a probe ranked by its platform-level mean count (x-axis, percentile), with $\log_{10}$(mean count + 1) on the y-axis. The dashed grey reference line indicates the 95$^{\text{th}}$ percentile of background signal. Target probes whose mean counts fall at or below this background threshold in MERSCOPE are highlighted and labeled. **B.** Spatial autocorrelation (Moran's I) for selected low-rank genes in MERSCOPE samples, comparing platforms within each gene facet. Each data point represent one sample, and boxplots summarize values across samples for Xenium (indigo, $n = 9$) and MERSCOPE (plum, $n = 7$).

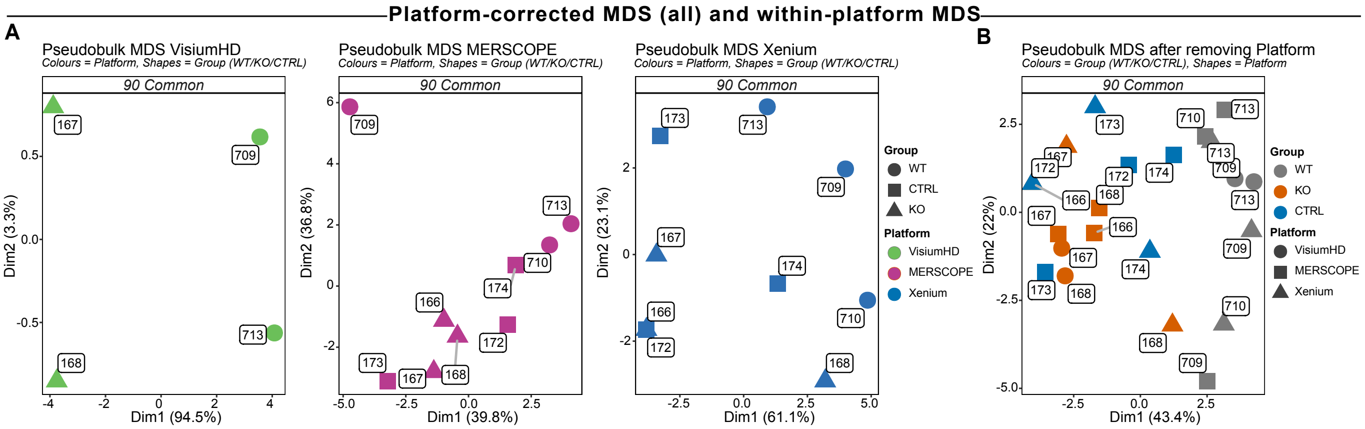

**Supplementary Fig. 5: Platform and genotype structure in pseudobulk spatial transcriptomics profiles.** **A.** Pseudobulk multidimensional scaling (MDS) of Visium HD, MERSCOPE and Xenium samples based on the shared 90-gene panel. Colours indicate platform and shapes indicate genotype (wild-type (WT), knockout (KO) and control (CTRL)). Sample identifiers are shown adjacent to each point. Axis labels indicate the percentage of variance explained by each dimension. **B.** Pseudobulk MDS of all samples following adjustment for platform effects using `limma::removeBatchEffect`[2]. Colors indicate genotype and shapes indicate platform.

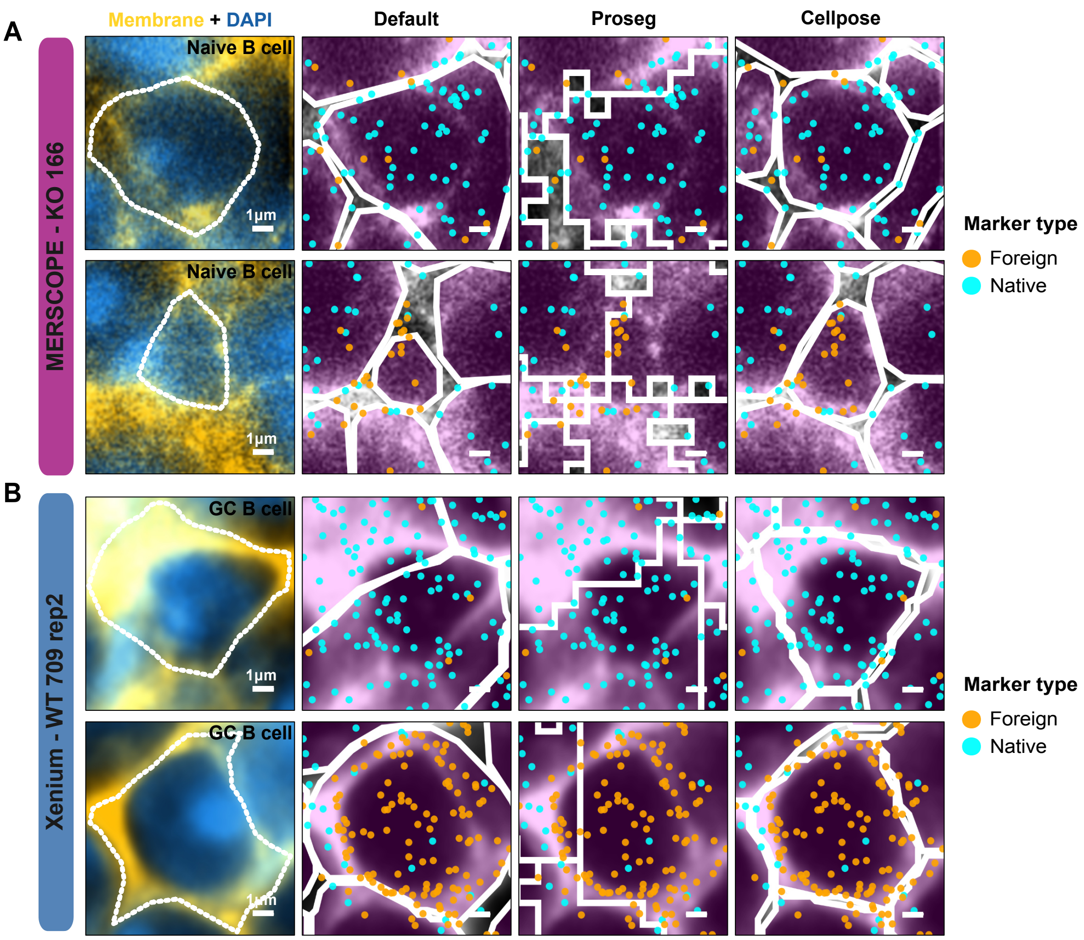

**Supplementary Fig. 6: Segmentation-dependent transcript assignment at cell boundaries in imaging-based spatial transcriptomics. A.** Representative naïve B cells from a MERSCOPE KO166 sample showing membrane and DAPI staining used for segmentation (left) and transcript assignment under three segmentation strategies (default pipeline, Proseg and Cellpose). White outlines indicate inferred cell boundaries. Detected transcripts are colored by marker class, distinguishing native cell-type markers (cyan) from foreign markers inconsistent with the annotated cell identity (orange). **B.** Representative germinal center B cells from a Xenium WT709 replicate analyzed using the same segmentation approaches, showing differences in transcript assignment at cell boundaries across segmentation strategies. Scale bars, 1 µm.

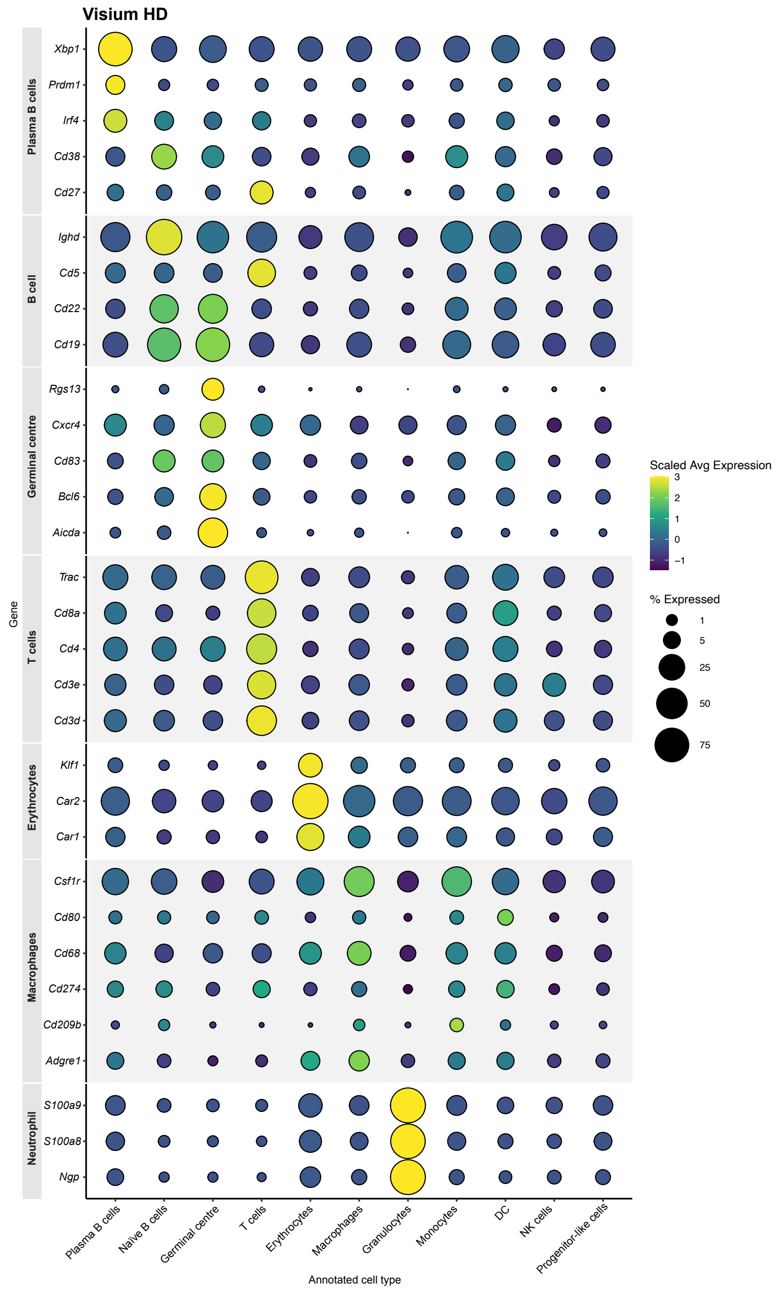

**Supplementary Fig. 7**: **Cell-type marker gene expression across annotated populations in Visium HD.** Dot plot of marker gene expression for cell types annotated using RCTD [3] in the Visium HD dataset. Dot color indicates scaled average expression and dot size indicates the percentage of bins in which each gene is detected. Marker genes include *Xbp1*, *Prdm1*, *Irf4*, *Cd38* and *Cd27* (plasma B cells); *Ighd*, *Cd5*, *Cd22* and *Cd19* (naïve B cells); *Rgs13*, *Cxcr4*, *Cd83*, *Bcl6* and *Aicda* (germinal center B cells); *Trac*, *Cd8a*, *Cd4*, *Cd3e* and *Cd3d* (T cells); *Klf1*, *Car1* and *Car2* (erythrocytes); *Csf1r*, *Cd80*, *Cd68*, *Cd274*, *Cd209b* and *Adgre1* (macrophages/monocytes); and *S100a9*, *S100a8* and *Ngp* (neutrophils).

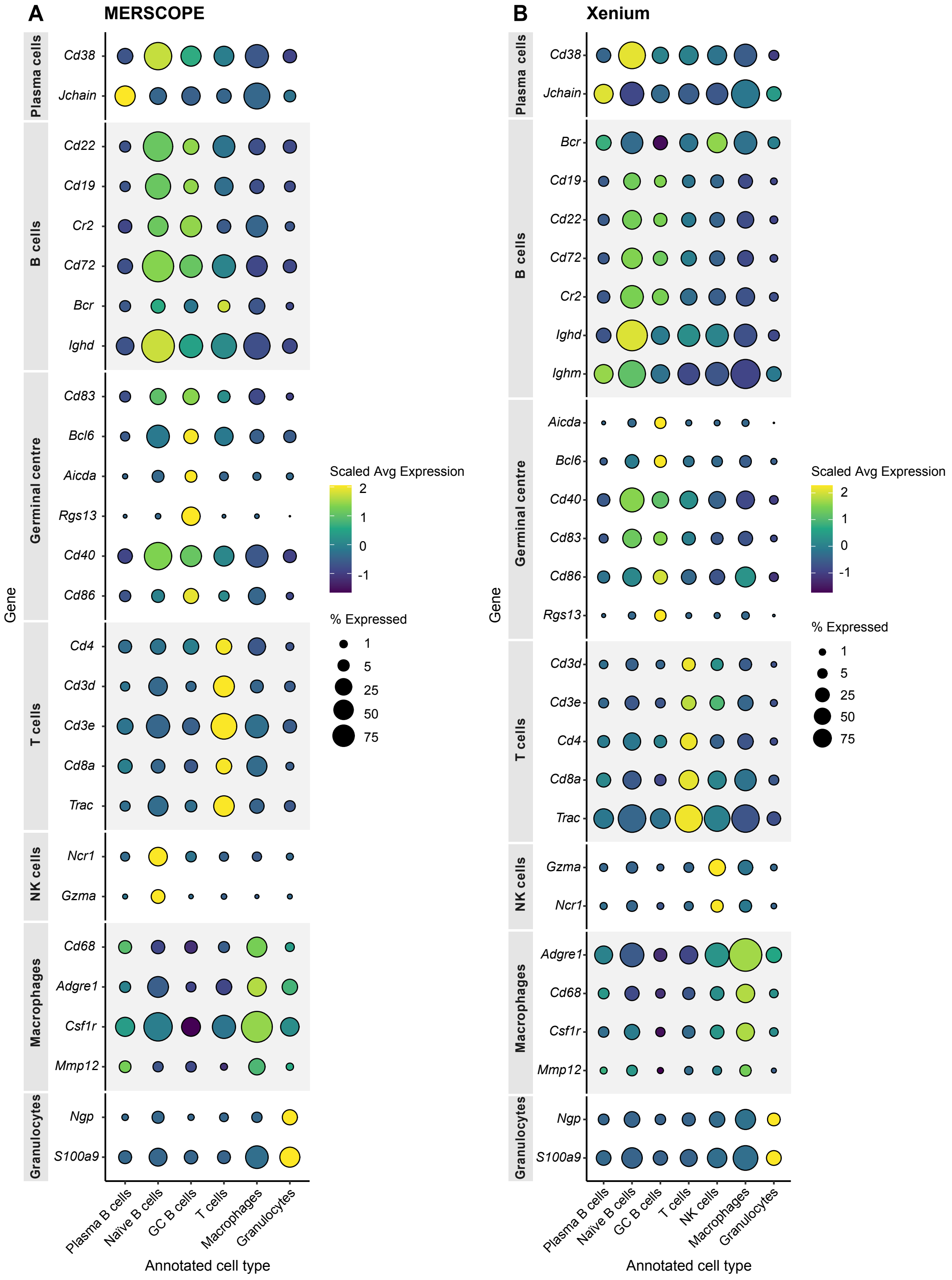

**Supplementary Fig.** 8: **Cell-type marker gene expression across annotated populations in MERSCOPE and Xenium.** Dot plots showing expression of representative marker genes across annotated cell populations for MERSCOPE (**A**) and Xenium (**B**). Dot colour indicates scaled average expression and dot size indicates the percentage of cells in which each gene is detected. Marker genes include *Jchain* (plasma B cells); *Cd19* and *Cr2* (B cells); *Cd83*, *Bcl6*, *Aicda* and *Rgs13* (germinal center B cells); *Cd4*, *Cd3d* and *Trac* (T cells); *Ncr1* (NK cells); *Adgre1* and *Csf1r* (macrophages); and *Ngp* and *S100a9* (granulocytes). NK, natural killer.

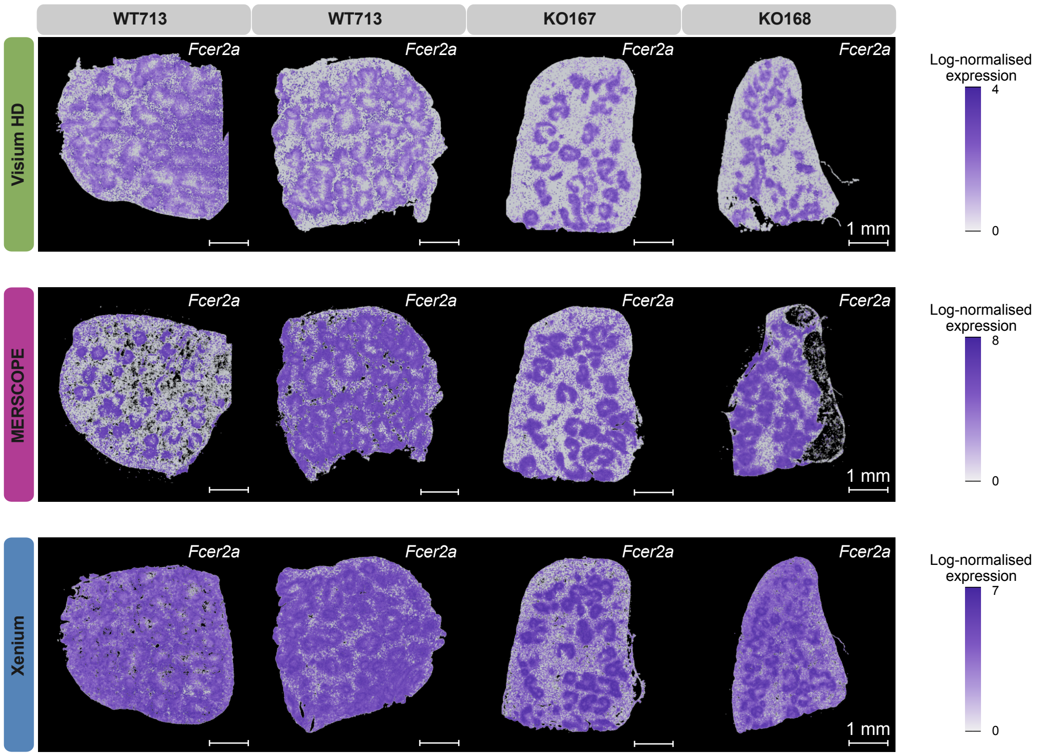

**Supplementary Fig.** 9: **Spatial expression of *Fcer2a* (Cd23) across spatial transcriptomics platforms.** Spatial maps showing log-normalised expression of *Fcer2a*, which encodes Cd23 and drives Cre recombinase expression in the Cd23-Cre model, across representative wild-type (WT) and Tbx21 conditional knockout (KO) spleen sections profiled using Visium HD, MERSCOPE and Xenium. Enrichment of *Fcer2a* signal within B-cell–rich follicular regions across platforms is consistent with expected Cd23-Cre activity in mature B-cell populations where deletion of the floxed *Tbx21* allele would occur. Scale bars, 1 mm.

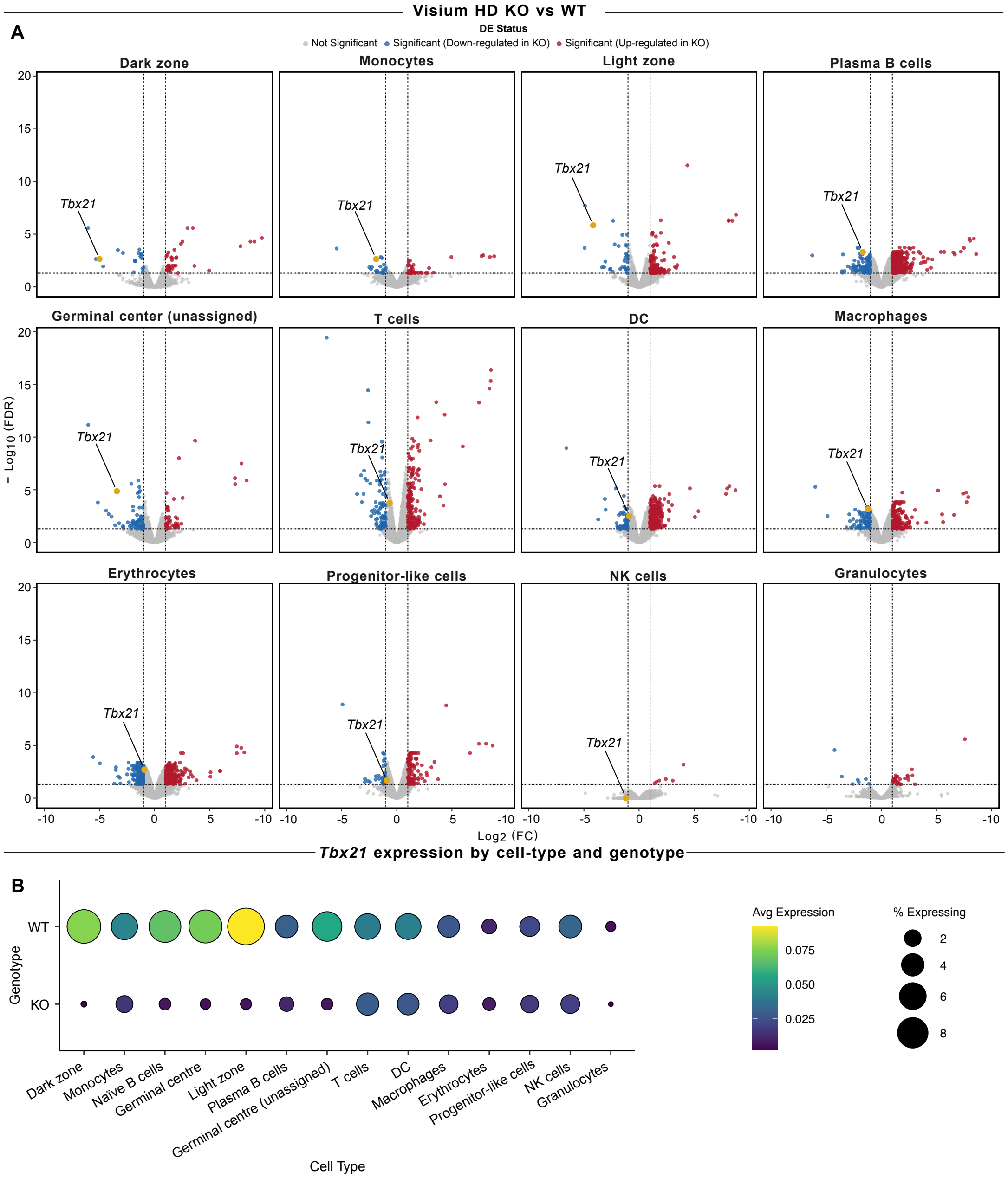

**Supplementary Fig.** 10: **Cell-type resolved differential expression of *Tbx21* in Visium HD knockout and wild-type samples. A.** Volcano plots showing pseudobulk differential expression between Tbx21fl/fl Cd23-Cre (KO) and wild-type (WT) samples for each annotated cell type in the Visium HD dataset. Differential expression was performed using the limma–voom [2,4] framework on pseudobulk counts aggregated by sample and cell type. Significance was defined as FDR $< 0.05$ and absolute $\log_2$FC $\geq 1$. **B.** Dot plot showing *Tbx21* expression across annotated cell types stratified by genotype. Dot colour indicates average normalized expression and dot size indicates the proportion of bins in which *Tbx21* transcripts were detected.

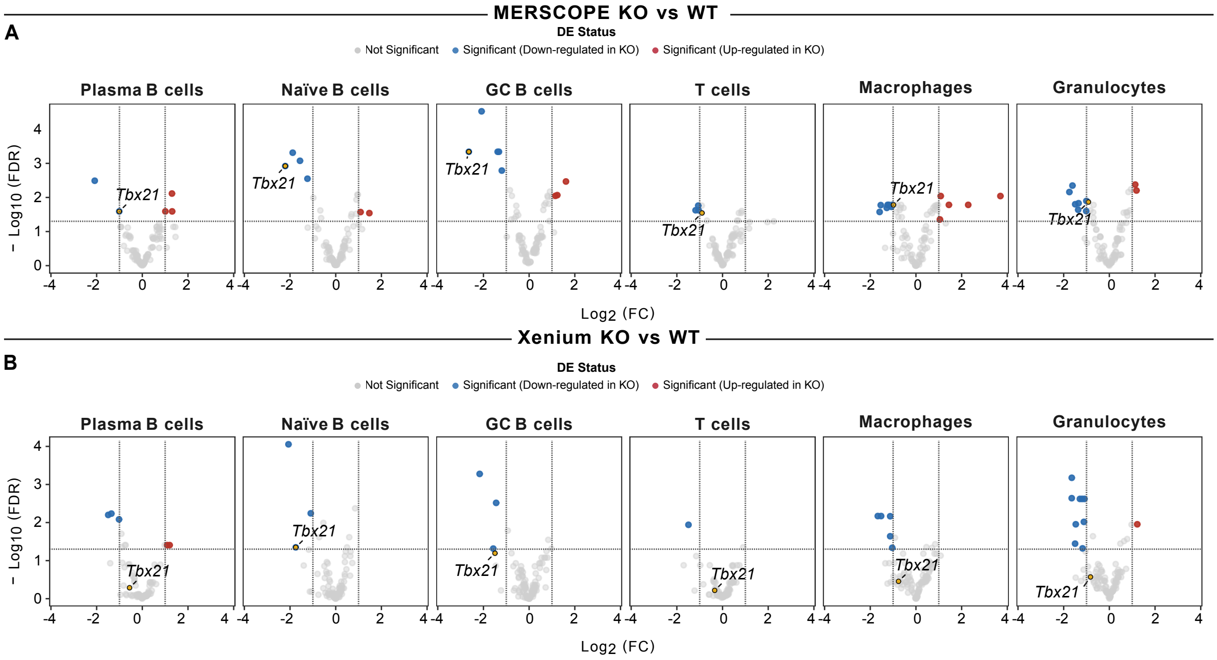

**Supplementary Fig.** 11: **Cell-type resolved differential expression of *Tbx21* in imaging-based spatial transcriptomics datasets. A.** Volcano plots showing pseudobulk differential expression between *Tbx21* conditional knockout (KO) and wild-type (WT) samples for each annotated cell type in the MERSCOPE dataset. Differential expression was performed using the limma–voom [2,4] framework on pseudobulk counts aggregated by sample and cell type. Significance was defined as FDR $< 0.05$ and absolute $\log_2$ fold change $\geq 1$. *Tbx21* is highlighted in each panel. **B.** Equivalent analysis for the Xenium dataset using the same pseudobulk differential expression framework and significance thresholds. *Tbx21* is highlighted in each panel.

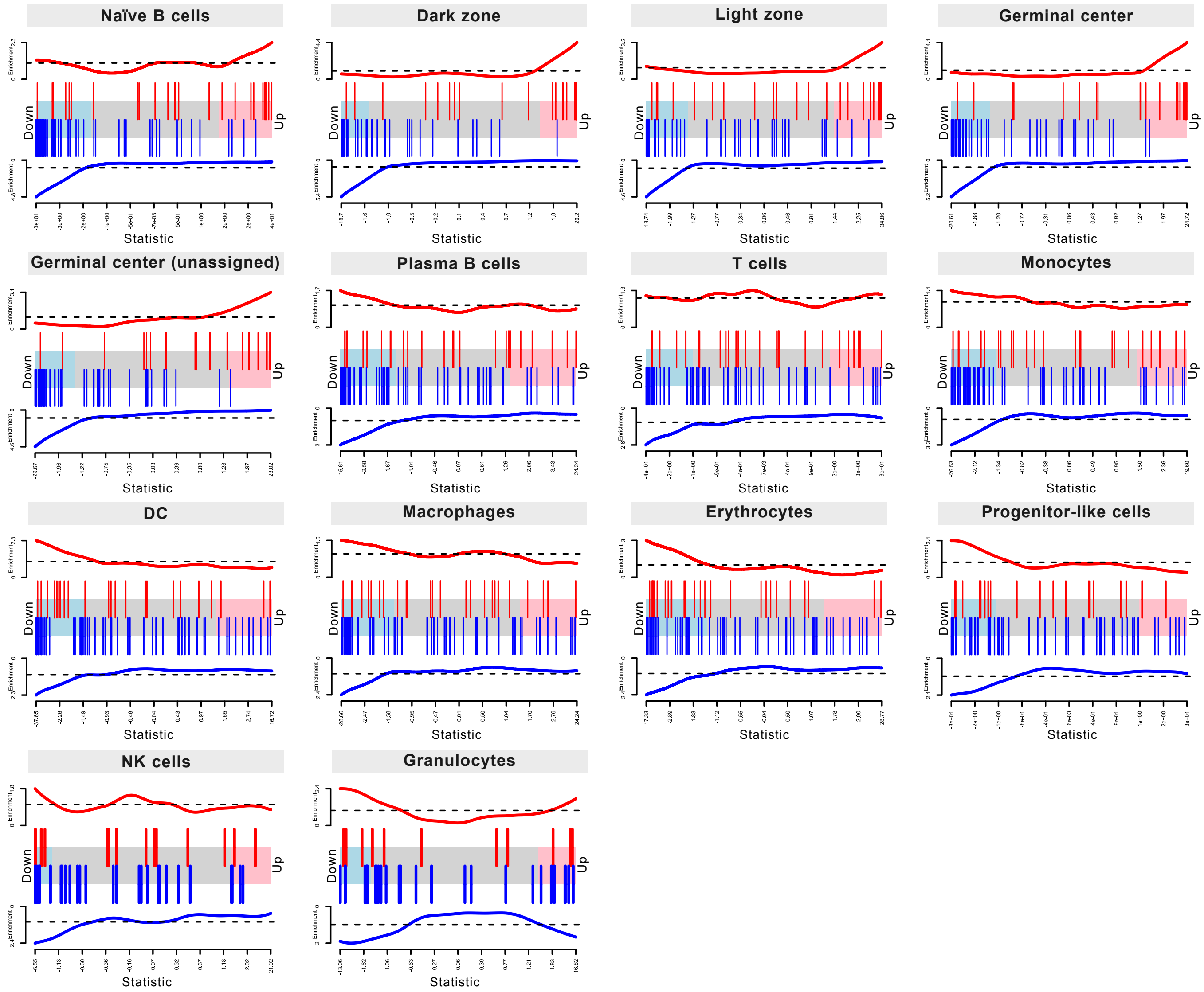

**Supplementary Fig.** 12: ***Tbx21*** **(Tbet) gene signature enrichment across cell types in Visium HD samples.** Rotation gene set testing (ROAST) barcode plots showing the distribution of genes from a published *Tbx21* germinal centre B-cell signature study by Ly et al.[(1)] across ranked pseudobulk differential expression statistics for each annotated cell type from *Tbx21* conditional knockout (KO) and wild-type (WT) samples. Genes are ranked from most downregulated in KO relative to WT (left) to most upregulated (right). Vertical bars indicate signature genes Coloured by direction (red upregulated, blue downregulated). Corresponding ROAST statistics are provided in **Supplementary Table 5**.

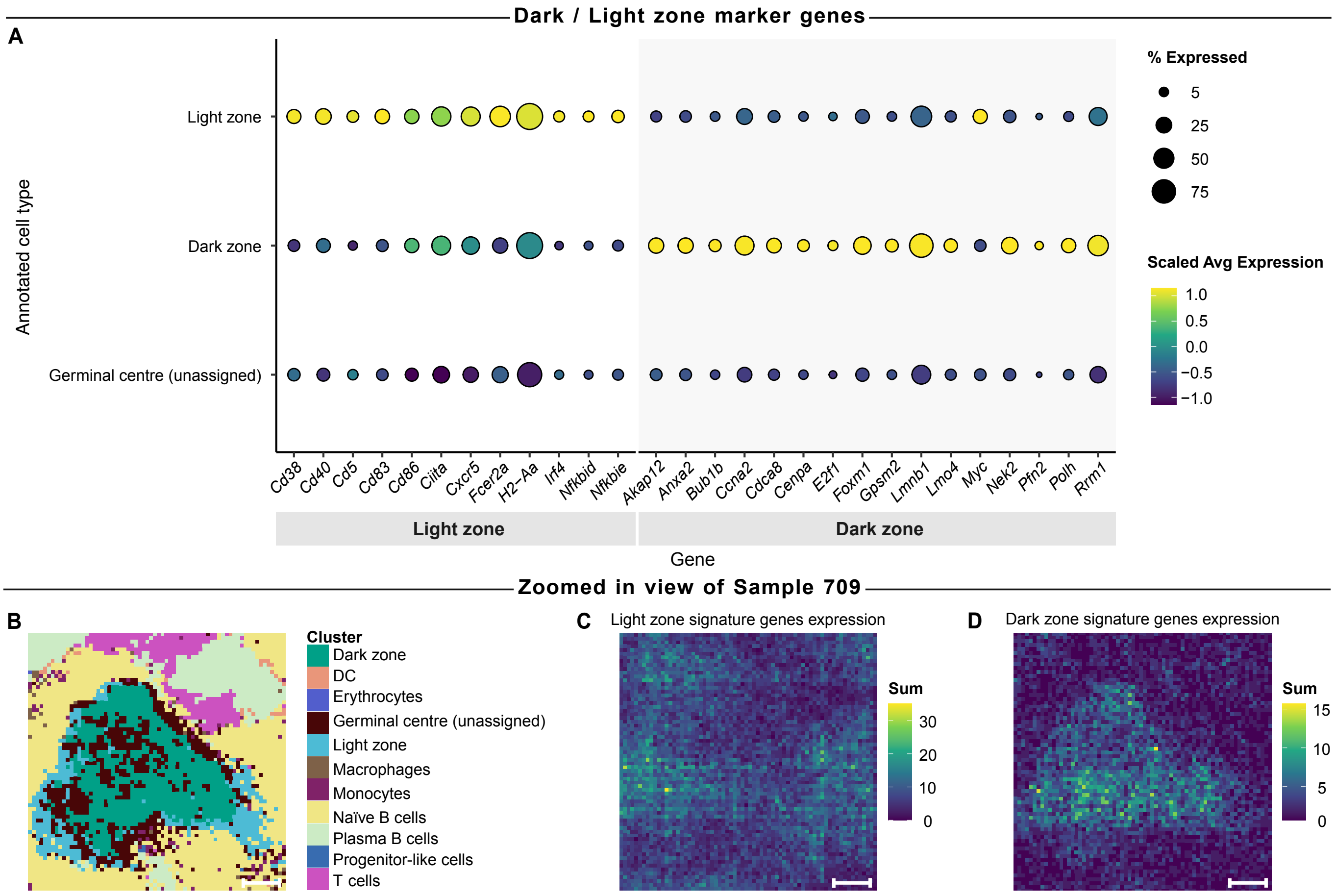

**Supplementary Fig.** 13: **Expression of germinal center dark and light zone marker genes and spatial zonation in Visium HD samples. A.** Dot plot showing expression of light zone (LZ) and dark zone (DZ) marker genes across annotated germinal center cell populations. Dot size indicates the percentage of cells expressing each gene and color indicates scaled average expression. **B.** Spatial map of annotated cell types in a representative Visium HD sample (WT709), highlighting DZ and LZ regions within the germinal center. Scale bar, 20 µm. **C-D.** Spatial heatmap showing the summed expression of LZ (**C**) and DZ (**D**) signature genes.

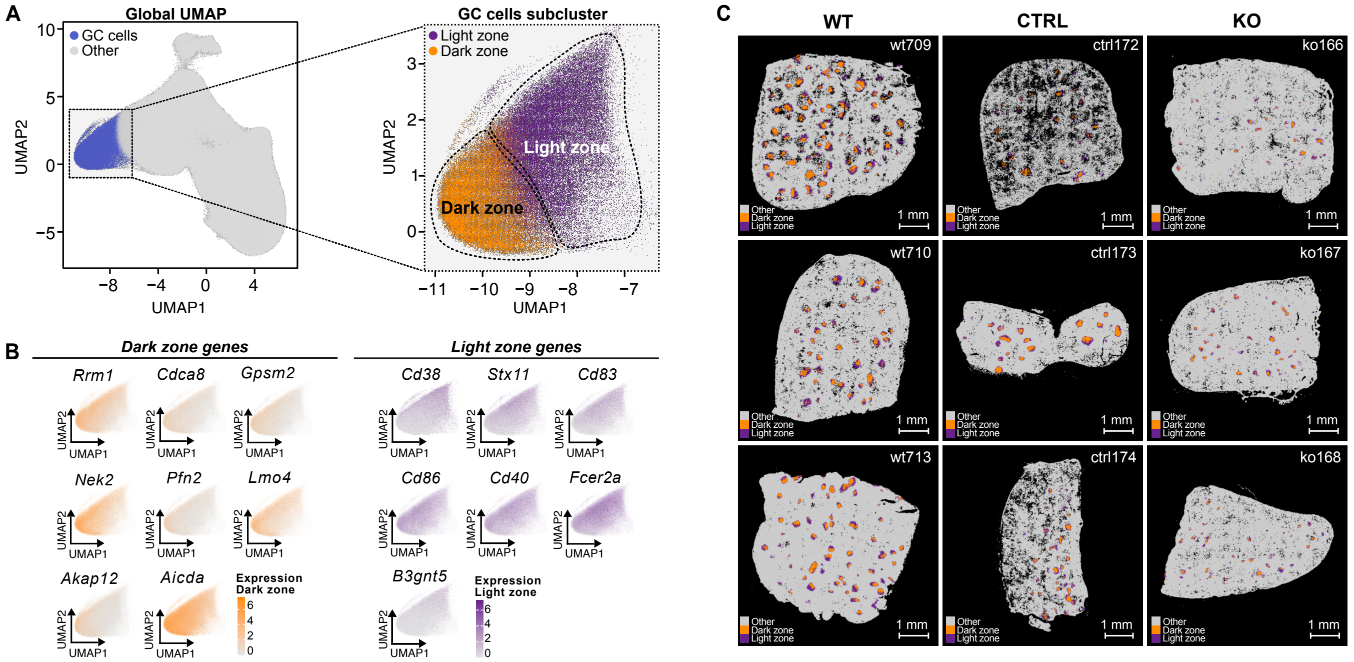

**Supplementary Fig.** 14: **Identification and spatial mapping of germinal center dark and light zones in Xenium samples. A.** Uniform Manifold Approximation and Projection (UMAP) of the integrated dataset highlighting germinal center cells among all other cell types (grey), with inset showing subclustering into dark zone (orange) and light zone (purple) populations. **B.** UMAP feature plots showing expression of dark zone (DZ) and light zone (LZ) marker genes used to define GC zonation states. **C.** Spatial distribution of dark and light zone cells across representative wild-type (WT), control (CTRL), and *Tbx21* conditional knockout (KO) spleen sections from the Xenium dataset. Scale bars, 1 mm.

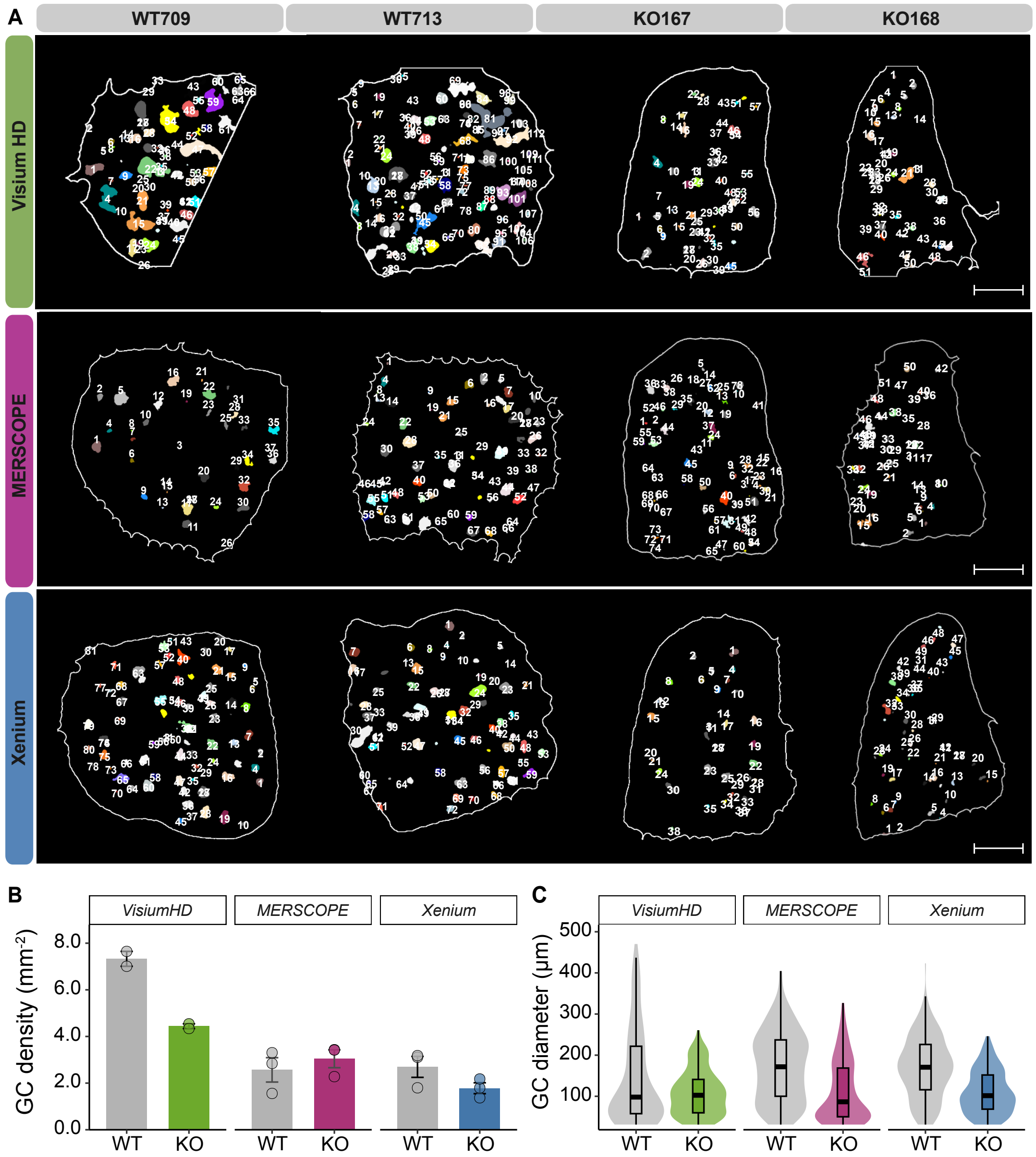

**Supplementary Fig.** 15: **Germinal center identification and morphological comparison across spatial transcriptomics platforms. A.** Spatial maps showing identified germinal centres (GCs) in wild-type (WT709, WT713) and *Tbx21* conditional knockout (KO167, KO168) spleen samples profiled with Visium HD, MERSCOPE, and Xenium. Numbers indicate individual GCs identified based on GC B cell and zonal annotations. Scale bars, 1 mm. **B.** Bar plot of GC density (GCs per $mm^2$) across WT and KO samples for each platform. Points indicate individual sample means. **C.** Distribution of GC diameters across platforms and genotypes. Violin plots show GC diameter distributions with boxplots indicating median and interquartile range. Sample sizes were Visium HD ($n = 2$ WT, $n = 2$ KO), MERSCOPE ($n = 3$ WT, $n = 3$ KO), and Xenium ($n = 3$ WT, $n = 3$ KO).

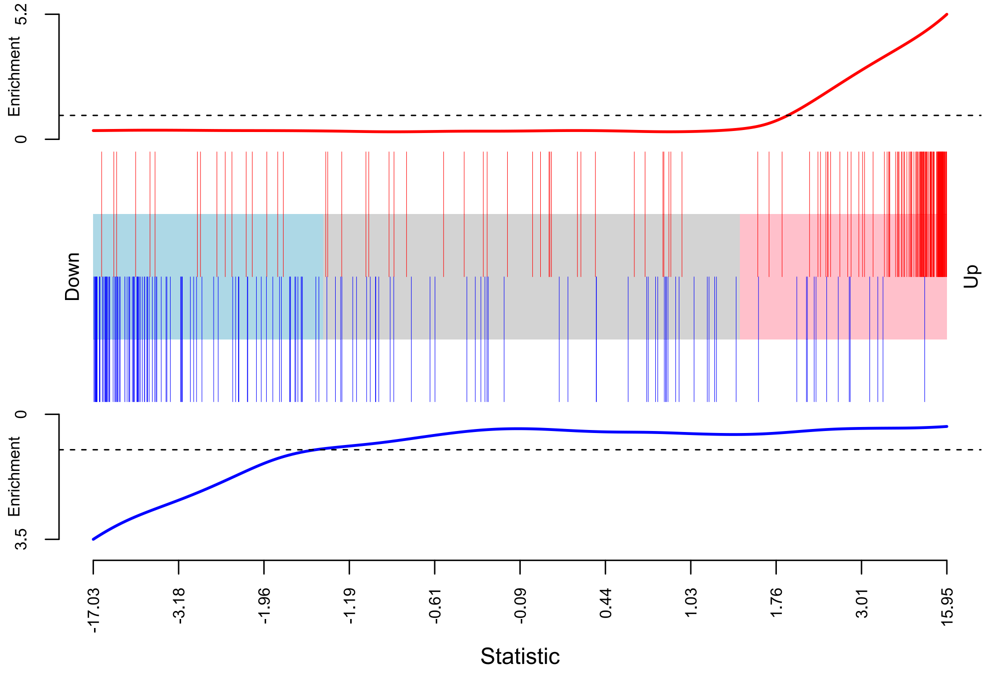

**Supplementary Fig.** 16: **Dark and light zone gene signature enrichment in Visium HD samples.** Rotation gene set testing (ROAST) barcode plot showing the distribution of genes from a published germinal center (GC) dark zone (DZ) versus light zone (LZ) gene signature study by Victora et al.[5] across ranked pseudobulk differential expression statistics from Visium HD wild-type samples. Genes are ranked from most downregulated in dark zone relative to light zone (left) to most upregulated (right). Vertical bars indicate signature genes colored by direction (red upregulated, blue downregulated). Corresponding ROAST test, $P = 0.007$ with 330 genes in common and an active proportion of 0.68.

#### Supplementary Information

**Supplementary Table.** 1: ***SpatialBench* sample metadata and processing information.** Summary of all samples processed across spatial transcriptomics platforms and sequencing assays included in this study. The table reports slide and batch identifiers, sample IDs, instrument software versions, inclusion status in the final analyses, and notes describing technical artifacts or quality issues encountered during processing. This table provides a complete record of all experimental attempts, including samples excluded from downstream analyses. The full table is provided in the Supplementary Data files as Supplementary Table 1.

**Supplementary Table.** 2: **Gene panel composition for MERSCOPE and Xenium assays.** List of genes included in the targeted spatial transcriptomics panels used for MERSCOPE and Xenium experiments in this study. The MERSCOPE panel comprises 91 genes and the Xenium panel comprises 100 genes. The full table is available in the Supplementary Data files as Supplementary Table 2.

**Supplementary Table. 3**: **Cellpose segmentation parameters for imaging-based spatial transcriptomics data.** Summary of model training, input image preprocessing, and inference settings used for Cellpose-based segmentation of MERSCOPE and Xenium images. Models were fine-tuned using a human-in-the-loop strategy starting from the pretrained *cyto2* model[(6,7)]. The table reports training dataset size, image preprocessing steps, augmentation strategies, and inference parameters used to generate cell segmentation masks for downstream transcript assignment and cell-level analyses.

| **Parameter** | **MERSCOPE** | **Xenium** |
|---|---|---|
| **Model and training** | | |
| Cellpose version | v2.3.2 | v2.3.2 |
| Training strategy | Human-in-the-loop fine-tuning | Human-in-the-loop fine-tuning |
| Initial model | Pretrained cyto2 | Pretrained cyto2 |
| Training images (n) | 78 | 89 |
| Testing images (n) | 14 | 17 |
| Epochs | 100 | 100 |
| Learning rate | 0.1 | 0.1 |
| Weight decay | $1 \times 10^{-4}$ | $1 \times 10^{-4}$ |
| **Input images and preprocessing** | | |
| Input channels | DAPI + composite greyscale (Cellbound2, Cellbound3, DAPI) | DAPI + composite greyscale (Cell Segmentation Stain) |
| Imaging plane used | $z = 3$ | $z = 3$ |
| Pixel size ($\mu$m) | 0.108 | 0.2125 |
| Image preprocessing | CLAHE normalisation (clip limit = 0.01) | CLAHE normalisation (clip limit = 0.01) |
| Data augmentation | Gaussian noise; blur; colour jitter | Gaussian noise; blur; colour jitter |
| **Inference and post-processing** | | |
| Inference software | `vpt-plugin-cellpose2` | QuPath v0.5.1 + BIOP Cellpose extension v0.9.6 |
| Cell diameter (px) | 66.94 | 33.88 |
| Flow threshold | 0.95 | 0.95 |
| Cell probability threshold | $-5.5$ | $-5.5$ |
| Output format | Cell-by-gene matrix regenerated via VPT | GeoJSON boundaries imported into Xenium Ranger v3.1 |

**Supplementary Table. 4: Differential expression results for Visium HD knockout versus wild-type across annotated cell types.** Gene-level results from pseudobulk differential expression analysis comparing KO and WT samples for each annotated cell type in the Visium HD dataset using the limma–voom[2,4] framework. Tables report estimated $\log_2$ fold changes and associated statistical measures for each gene. Details of the pseudobulk analysis and statistical modeling are described in the *Methods*. The complete results are provided as Supplementary Data files as Supplementary Table 4.

**Supplementary Table. 5: ROAST gene set concordance with Ly et al.[1] using Visium HD KO versus WT differential expression results.** Rotation gene set testing (ROAST) was applied to pseudobulk differential expression results from the Visium HD knockout versus wild-type comparison for each annotated cell type. Enrichment was assessed against the Tbet–associated germinal center gene program reported by Ly et al.[1]. The active proportion indicates the fraction of genes contributing to the directional enrichment signal, and the P-value represents the statistical significance of concordance with the reference gene set.

| Cell Type | Active Proportion | P-Value | Number of Genes |
|---|---|---|---|
| Dark zone | 0.640 | 0.000670 | 86 |
| Monocytes | 0.412 | 0.000795 | 85 |
| Naive B cells | 0.602 | 0.000810 | 88 |
| Germinal centre | 0.652 | 0.001040 | 89 |
| Light zone | 0.593 | 0.001290 | 86 |
| Plasma B cells | 0.437 | 0.001630 | 87 |
| Germinal centre (unassigned) | 0.581 | 0.002000 | 86 |
| T cells | 0.333 | 0.004500 | 84 |
| DC | 0.288 | 0.008700 | 80 |
| Macrophages | 0.313 | 0.011100 | 83 |
| Erythrocytes | 0.356 | 0.033200 | 90 |
| Stem cells | 0.270 | 0.063300 | 74 |
| NK cells | 0.150 | 0.066100 | 40 |
| Granulocytes | 0.256 | 0.199000 | 39 |

**Supplementary Table. 6: Differential expression results between germinal center dark and light zones in Visium HD wild-type samples.** Gene-level results from pseudobulk differential expression analysis comparing dark zone and light zone germinal center regions in Visium HD wild-type samples using the limma-voom[2,4] framework. Tables report estimated $\log_2$ fold changes and associated statistical measures for each gene. Details of the pseudobulk analysis and statistical modeling are described in the *Methods*. The complete results are provided in the Supplementary Data files as Supplementary Table 6.
